# Designing urban green-space heterogeneity across urbanisation gradients to maintain avian richness

**DOI:** 10.64898/2026.02.12.705555

**Authors:** Jie Hu, Danique van Os, Joeri Morpurgo, Michiel P. Veldhuis, Roy P. Remme, Geert de Snoo, Yali Si

## Abstract

Urban expansion drives land cover change and habitat simplification, contributing to biodiversity loss. Urban green spaces can mitigate these impacts, but their effectiveness depends on its configuration and implementation. Here, we examine how three complementary dimensions of environmental heterogeneity—plant species richness, habitat heterogeneity, and foliage-layer richness—shape bird richness along an urbanisation gradient in the Netherlands. Using bird and plant occurrence data, LiDAR-derived vegetation structure, and land-use data, we fitted generalized additive models at three spatial scales (100, 200, and 300 m) to assess how these relationships vary across the urbanisation gradient. Plant species richness showed the strongest and consistent positive effect on bird richness, disregarding urbanization intensities. Habitat heterogeneity showed most pronounced positive effects at intermediate levels of urbanisation. In contrast, foliage-layer richness had weak associations with bird richness across urbanization intensities. Together, these results demonstrate that sustaining urban bird diversity requires urbanisation-intensity–dependent design of green-space heterogeneity. Increasing plant richness is generally recommended across urbanization intensities. Increasing habitat heterogeneity is more effective at intermediate levels of urbanisation and appears less suitable in highly urbanised contexts. Beyond simply expanding green space area or their spatial complexity alone, urban planning should focus on the thoughtful design of different types of environmental heterogeneity. This includes city-wide species-rich planting and structurally diverse habitat mosaics in mid-density areas to sustain urban bird diversity.

## 1. Introduction

Urban expansion drives land-cover change, leading to biodiversity loss (Casanelles-Abella et al., 2025; Öztürk et al., 2022). In response, cities increasingly seek to reconcile human development with biodiversity goals (Casanelles-Abella et al., 2025; Goddard et al., 2010; Sockhill et al., 2025). Policy frameworks, including the EU Nature Restoration Law (EU, 2024), CBD (CBD, 2022) and the EU Birds Directive (EU, 2009), are now recognizing urban green-space design and management as a tool towards bending the curve of biodiversity loss. Urban green spaces offer opportunities to promote biodiversity by providing novel ecological habitat and connectivity which are known to support species populations already persisting or thriving in cities (Kang et al., 2015; Morpurgo et al., 2023; Oropeza-Sánchez et al., 2025; Rudd et al., 2002). However, the degree to which they deliver these benefits varies with design and management (Melles et al., 2003; Morpurgo et al., 2025). Hence a core question is how to design and manage urban green spaces to optimize their contribution to biodiversity (Goddard et al., 2010; Kang et al., 2015; Sockhill et al., 2025; Von Thaden et al., 2021).

Environmental heterogeneity, defined as variation in environmental conditions, is a key mechanism through which urban green spaces can support biodiversity, by providing diverse niche opportunities, reducing stress through shelter and offering refugia from disturbance (Stein et al., 2014; Tews et al., 2004; Threlfall et al., 2017). Environmental heterogeneity is commonly partitioned into five components: two biotic components (habitat and vegetation) and three abiotic components (climate, topography, and soil) (Stein et al., 2014; Stein & Kreft, 2015). In urban systems, habitat heterogeneity (also referred to as land cover heterogeneity or habitat diversity) and vegetation heterogeneity (e.g., plant diversity, plant complex) are key contributors to biodiversity, as they directly shape the availability and diversity of ecological resources (Schütz & Schulze, 2015; Sultana et al., 2021; Threlfall et al., 2017) while also influencing visual qualities such as colour and texture (de Val et al., 2006; Dronova, 2017). Importantly, these biotic components are also among the elements most directly and rapidly modified through human management, making them particularly relevant for urban planning and design (Fernández-Juricic, 2000; Souza et al., 2019). For example, bird habitat diversity can be achieved through the composition and spatial configuration of public parks and private gardens (Moudrý et al., 2021; Souza et al., 2019); when well designed, such habitat mosaics can support nesting and foraging opportunities while also increasing perceived visual richness for residents (de Val et al., 2006). Consequently, using environmental heterogeneity as a planning tool to conserve urban biodiversity requires a robust understanding of the relationship between environmental heterogeneity, particularly in habitat and vegetation, and biodiversity patterns in cities.

Environmental heterogeneity–biodiversity relationships might vary along the urbanisation gradient (i.e. from low to high urbanization and population density) due to an area–heterogeneity trade-off (Ben-Hur & Kadmon, 2020; Chiron et al., 2024). Increasing heterogeneity can enhance the diversity of resources and environmental conditions at the site level, yet it may simultaneously reduce the amount of suitable habitat available to any given species by subdividing space into smaller patches (Kang et al., 2015; Oropeza-Sánchez et al., 2025; Watson et al., 2005; White et al., 2005). In small and fragmented urban green spaces, this trade-off can translate into microhabitat fragmentation, stronger edge effects, and reduced local population sizes, thereby weakening or even reversing heterogeneity benefits (Havlíček et al., 2021; Schütz & Schulze, 2015; Suarez-Rubio et al., 2018). This explains why some studies report positive effects of environmental heterogeneity on biodiversity in urban landscapes (Peng et al., 2024; Sockhill et al., 2025; Threlfall et al., 2017), others find negative or non-significant relationships (Schütz & Schulze, 2015; Suarez-Rubio et al., 2018). Although previous studies have examined the effects of vertical vegetation heterogeneity (Bradbury et al., 2005; Sockhill et al., 2025), plant species richness (da Silva et al., 2021) and habitat heterogeneity (Melo et al., 2022; Sultana et al., 2021), the change of the relationships across varying levels of urbanisation intensities has rarely been investigated. This distinction is especially relevant for urban planning, where intervention strategies need to be tailored to the degree of built-up intensity.

To address this gap, it is necessary to focus on an indicator species that responds sensitively to variation in habitat and vegetation structure and is relevant to urban planning and management decisions. Birds are particularly informative indicators of environmental heterogeneity–biodiversity relationships, as they are data-rich (La Sorte et al., 2018), differ markedly in dispersal ability and habitat preferences (Grinnell, 1917; Karr & Freemark, 1983) and show distributions closely associated with plant species composition and vertical vegetation structure (Donnelly & Marzluff, 2004; Lepczyk & Warren, 2012). Because bird species have a wide range of movement distances and space-use strategies (Tamburello et al., 2015) and bird communities integrate habitat conditions across multiple spatial scales (Odum & Kuenzler, 1955). This makes birds a sensitive lens for evaluating how heterogeneity-based design and management strategies influence biodiversity outcomes in urban landscapes (Food & Affairs, 2003; Paker et al., 2014). In the Netherlands, where approximately 16% of land is classified as urban, nearly one-third of breeding bird species have a disproportionately large share of their breeding pairs in urban areas (Snep et al., 2016), highlighting both the ecological relevance of cities for avian biodiversity and the value of birds as indicators for urban biodiversity planning.

This study aims to assess how environmental heterogeneity-based approaches can be used to maintain biodiversity along gradients of built-up intensity. We first characterised spatial patterns of key biotic components of environmental heterogeneity, plant species richness, habitat heterogeneity, and foliage-layer richness, along the gradient of built-up cover in Dutch urban landscapes. We then tested how each heterogeneity component influences bird species richness across urbanization intensities. Our findings provide evidence for the context dependence of environmental heterogeneity management as a strategy for enhancing urban biodiversity and offer spatially explicit insights to support landscape architects and urban planners in designing and managing green spaces that are both ecologically effective and context-sensitive.

## 2. Methods

### 2.1 Study area

The study area covers approximately 3,100 km^2^ in the urbanised western Netherlands and spans a polycentric metropolitan system centred on the country’s four largest cities, Amsterdam, The Hague, Rotterdam and Utrecht (Fig. 1). This region has been shaped by long-standing spatial planning aimed at managing growth across multiple urban centres (Kühn, 2003). It is characterised by high population density and a dense network of road and water infrastructure (Geurs & Rietveld, 2006). Together, these features create strong and continuous variation in built-up cover, from dense urban cores to peri-urban neighbourhoods, providing an ideal urbanisation gradient for testing how environmental heterogeneity relates to bird species richness. Because our focus is on urban residential and mixed-use fabrics, agricultural landscapes were excluded from the analysis (Kühn, 2003). Large infrastructure and industrial complexes (e.g., Schiphol Airport and parts of the Port of Rotterdam) were also excluded for the same reason.

**Figure 1:**
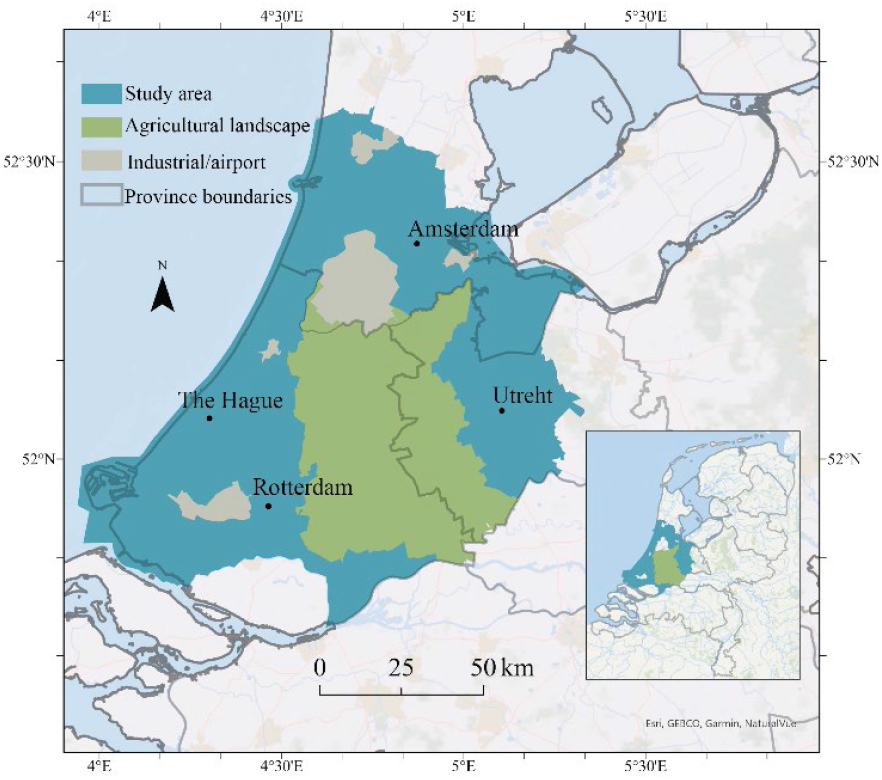
Study area within the Randstad region (western Netherlands; approximately 3,100 km^2^). The study area used in this analysis is shown in blue and spans the metropolitan region around Amsterdam, Rotterdam, The Hague and Utrecht. Agricultural landscapes (green), and major infrastructure/industrial complexes (grey; including Schiphol Airport and parts of the Port of Rotterdam) were excluded from analysis. Province boundaries are shown as grey outlines. The inset map indicates the location of the study area within the Netherlands. Basemap: Esri World Ocean.

### 2.2 Data

Bird observation data used to calculate bird richness were obtained from the Dutch National Database of Flora and Fauna (NDFF; https://www.ndff.nl/). Data quality is ensured through automated rule-based checks and expert validation of flagged records, providing high reliability in species, spatial and temporal information. For this study, we utilized data only from 2020, which is encompassing observations of 350 bird species within the study area.

Vascular plant species data were also obtained from the NDFF to characterise plant richness. The dataset documents the occurrence of 1,342 plant species across the study area. Unlike the bird data, which are point-based observations, plant records were collected through plot-based surveys, with each species occurrence assigned to the central coordinate of its survey plot. We selected all records collected in plots of 50m^2^ or smaller for each grid cell in the same year (2020), and summed the unique plant species to calculate plant species richness.

Vegetation foliage layers (Vegetation vertical structure) data were obtained from the dataset country-wide data of ecosystem structure derived from the third Dutch airborne laser scanning survey (https://zenodo.org/records/6421381), with a spatial resolution of 10 meters (Kissling et al., 2023). This dataset was generated using the Actueel Hoogtebestand Nederland (AHN; https://www.ahn.nl/, Version 3) and includes 25 ecosystem structure metrics. Each metric was calculated from LiDAR point cloud data within 10 × 10 meter grid cells (Kissling et al., 2022, 2023). We classified vegetation height into four foliage layers: grass (< 1 m), shrub (1–3 m), tree (3–20 m) and tall tree (> 20 m). To ensure that LiDAR data reflected non-built surfaces, points overlapping with building footprints were removed using the 2021 Dutch cadastre’s Basisregistratie Adressen en Gebouwen dataset (BAG; https://bagviewer.kadaster.nl/).

Habitat types data used to calculate habitat heterogeneity were derived from the Landelijk Grondgebruiksbestand Nederland (LGN; https://lgn.nl/, Version 2020) at a spatial resolution of 5 meters. The dataset includes 48 land-use categories and we reclassified these classes into 23 ecologically meaningful habitat types relevant for birds (see Appendix, Table S1).

### 2.3 Environmental heterogeneity

The spatial scales used in this study were chosen based on the Netherlands bird species’ typical home-range radius, spanning from 0.1 to 15 km (Appendix, Table S2), but 49% are smaller than 0.2 km (Tamburello et al., 2015). Therefore, we focused on three fine-grained spatial scales100m, 200m, 300m (grid cell sizes of 100 × 100 m, 200 × 200 m, and 300 × 300 m). Bird species richness at each scale was quantified as the number of species recorded within each grid cell. For the statistical modelling, we used a log-transformed response, log(1 + bird species richness), to stabilise variance and reduce the influence of extreme values.

We considered three components of environmental heterogeneity: habitat heterogeneity, plant richness and foliage-layer richness. Habitat heterogeneity within each grid cell was quantified using the Shannon diversity index. Plant richness was measured as the number of unique plant species recorded per grid cell; for modelling we used log-transformed plant richness, log (1 + plant species richness), to reduce right skew and the influence of extreme values and to better capture diminishing marginal changes in richness when estimating smooth and interaction terms. Foliage-layer richness was quantified as the count of grass, shrub, tree and tall tree layers within a grid cell, representing the complexity of vertical vegetation structure.

Additionally, we used the LGN data to calculate the proportion of built-up cover as an indicator of urbanization. Built-up area was defined by aggregating anthropogenic land-use classes into a single category, including residential buildings (primary and secondary), suburban development, roads and railroads, bare ground in residential areas, and greenhouses (Appendix, Table S1). For each spatial scales, built-up cover (%) was calculated as the proportion of the cell area defined as built-up, and used to represent the degree of urbanization. Built-up cover was included as a continuous predictor in model fitting; categorisation was used only for result visualisation. For response-curve plots, we generated predictions at three representative built-up cover levels: low (10%), medium (50%), and high (80%), chosen to represent well-sampled portions of the urbanisation gradient, low (“rural”), medium (“peri-urban”) and high (“urban”).

### 2.4 Data analysis

Prior to modelling, we examined pairwise correlations among these environmental predictors and calculated variance inflation factors (VIFs) to assess collinearity. The included predictors correlations were < 0.4 and all VIFs < 2, and considered appropriate for regression analysis (Appendix, Table S3-S4). To investigate the relationship between environmental heterogeneity and bird richness along gradients of built-up intensity, we applied a Generalized Additive Model (GAM) including both main effects and interactions with built-up cover. GAMs from the *mgcv* package for R (Wood, 2025) extend traditional linear models by capturing both linear and non-linear trends through smoothing splines, which are smoothed curves fitted to the data points. This method enables the detection of non-linear relationships and reduce research input bias for choice of non-linear functions. To ensure unbiased parameter estimation and optimal smoothing, the model was estimated using the Restricted Maximum Likelihood (REML) method.

The general formula used to fit the data in our study is:

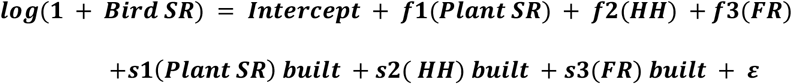

where *Bird SR* is bird species richness, *Plant SR* is log-transformed plant richness, *HH* is habitat heterogeneity, *FR* is foliage-layer richness, and built is built-up cover (%) representing the urbanisation gradient. The functions *f* are smooth main effects of each heterogeneity component, fitted using shrinkage thin plate regression splines in mgcv (bs = “*ts*”). The terms *Built×f* represent varying-coefficient smooths (implemented using by = *built*), allowing the shape and strength of each heterogeneity–richness relationship to change along the urbanisation gradient. The intercept represents the overall mean of *log(1 + Bird SR)* on the model scale because smooth terms in *mgcv* are centred to have zero mean contribution. The error term ε denotes residual variation. Model adequacy was evaluated using basis-dimension checks (Appendix, Table S5). To assess non-linear collinearity among smooth components, we quantified and reported concurvity metrics in the Appendix (Appendix, Table S6).

To assess the robustness of our results and to evaluate how the relationship between environmental heterogeneity and bird richness changes across spatial scale, we fitted three identical GAMs at spatial scales of 100m, 200m and 300m. All heterogeneity predictors and interactions were significant at all scales, and the shapes of the smooth functions were qualitatively similar (Appendix, Fig. S2-3). Model fit improved with increasing scale (deviance explained: 13% at 100 m, 21% at 200 m, 26% at 300 m; Appendix, Table S7), with very similar responses at 200 m and 300 m. Considering both model performance and the typical bird home-range sizes, we present the 200 m models in the main text and treat the 100 m and 300 m results as a sensitivity analysis (Appendix, Fig. S2-3, Table S7) (Davison et al., 2023).

## 3. Results

### 3.1 Environmental heterogeneity in urban landscapes

The study area contains non-built-up areas along the fringes and in larger natural areas, with higher built-up areas in the major cities (Fig. 2A). Plant richness (Fig. 2B, Fig 2E) spans the entire gradient and appears more spatially intermixed. Medium richness levels (20–60 species) are most common in areas with 30–60% built-up cover (Fig. 2B, Fig. 3A), but both very low (<20 species) and very high (>100 species) species occur in isolated patches across the landscape. Habitat heterogeneity (Fig. 2C, Fig 2E) presents a more uniform distribution across the landscape. Medium (0.67–1.20) to medium– high (1.78–2.22) habitat heterogeneity level are concentrated in areas with medium to high built-up density (>50%, Fig. 2C, Fig. 3B), while small pockets of very high heterogeneity are found in regions with minimal built-up area (<20%, Fig.2C, Fig. 3B). Foliage layer richness (Fig. 2D, Fig 2E) exhibits a clear spatial pattern: medium values (2–3) dominate the urban core, forming broad, cohesive clusters in areas with high built-up coverage (>70%, Fig. 2D, Fig.3C). In contrast, very low (=1) and very high (=4) foliage layer richness are primarily located along the urban fringe, appearing as belts or scattered fragments in regions where built-up cover is low (<30%, Fig. 2D, Fig. 3C).

**Figure 2.**
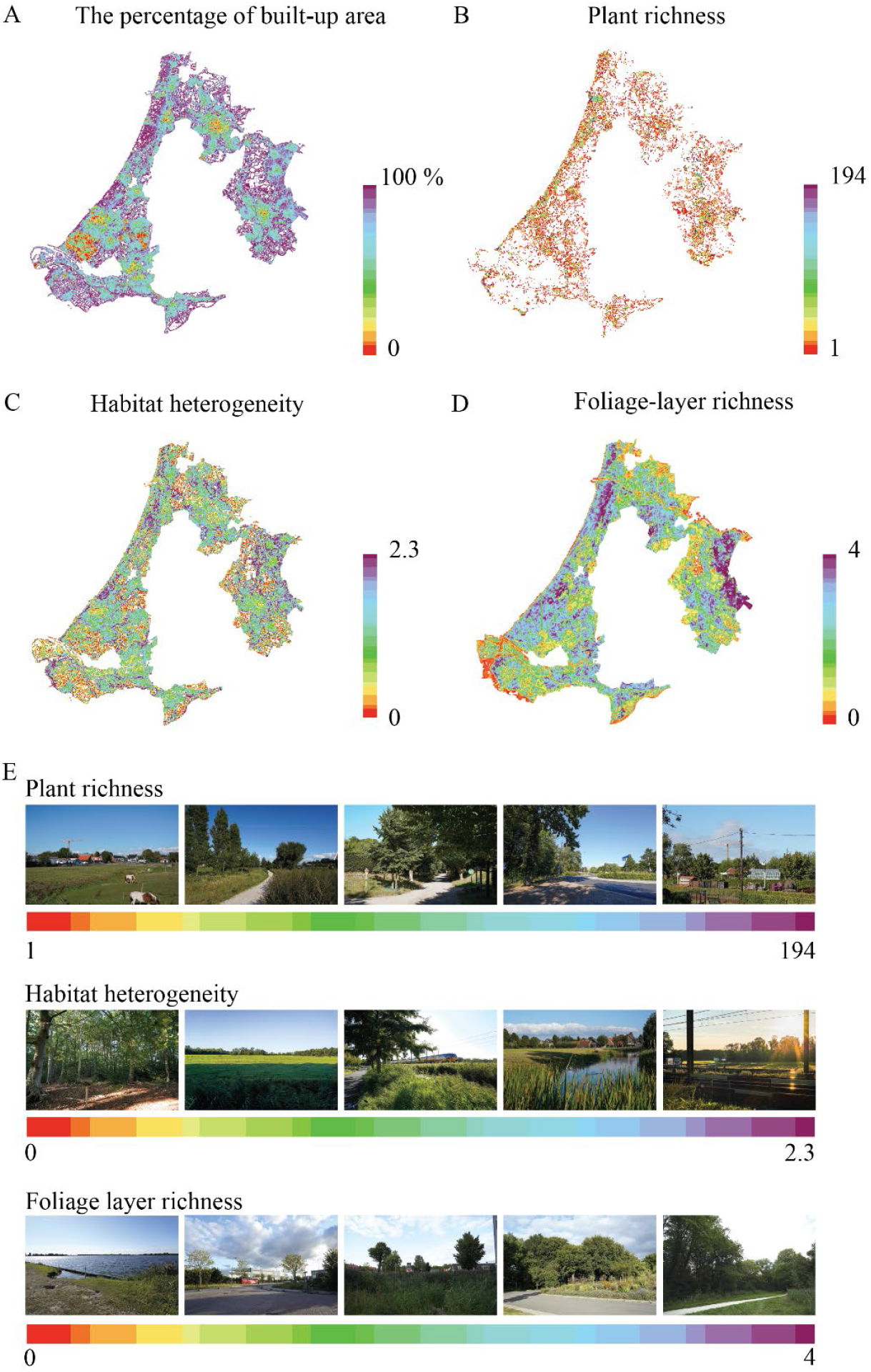
Environmental heterogeneity varies markedly across the built-up gradient, with plant species richness, habitat heterogeneity, and foliage-layer richness showing distinct spatial patterns across the urban landscape in the Netherlands. (A) Percentage of built-up area, (B) plant species richness, (C) habitat heterogeneity (Shannon index), and (D) foliage-layer richness, all mapped at 200-m resolution. Colour scales show raw values from low (yellow–red) to high (blue–purple): built-up cover 0–100%, plant richness 1–194 species, habitat heterogeneity 0–2.3, foliage-layer richness 0– 4 layers. (E) Photo strips provide a “zoom-in” view of typical local environments along the gradients in panels B–D; photographs are illustrative only and were not used in the analyses. Colour bars under each strip correspond to the mapped value ranges.

**Figure 3.**
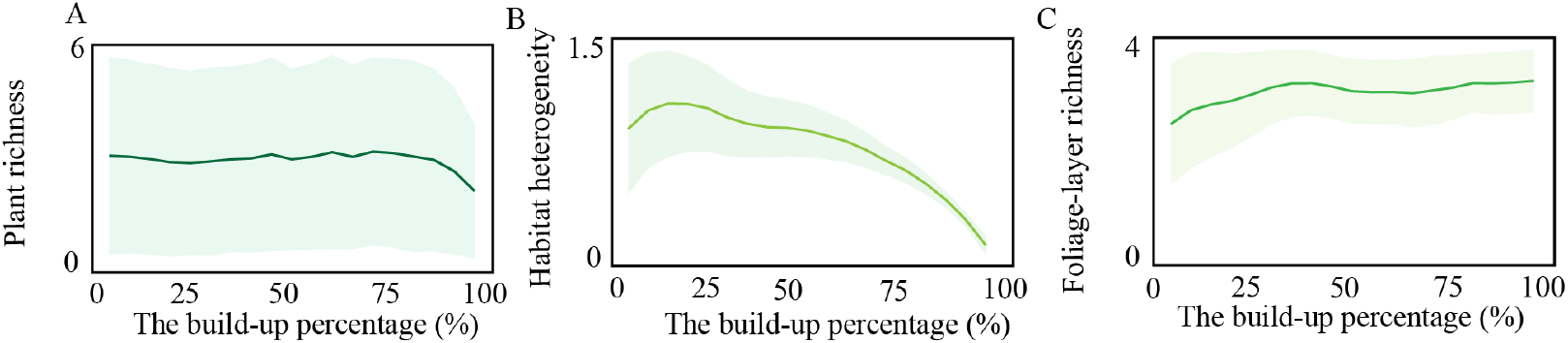
Plant species richness remains relatively stable across the built-up gradient, whereas habitat heterogeneity declines sharply with increasing urbanisation and foliage-layer richness shows a slight increase. Environmental heterogeneity across the built-up area gradient (0–100%). Lines show mean values and shaded bands show ± standard deviation SD, computed in 5% built-up bins and plotted at bin midpoints. Panels: (A) plant species richness, (B) habitat heterogeneity, and (C) foliage-layer richness. For visualization, plant species richness values outside the 2.5th and 97.5th percentiles were excluded to reduce the influence of extreme observations; all statistical analyses used the full dataset.

### 3.2 Bird richness driven by environmental heterogeneity across urbanization intensity

All three components of environmental heterogeneity were significantly associated with bird richness, and their effects varied along the urbanisation gradient (Table 1; Fig. 4). Overall, bird richness decreased with increasing built-up cover (Fig. 4). Plant species richness showed a strong and consistently positive association across the entire gradient, although the increase in bird richness was steepest in greener areas and became more gradual as built-up cover increased (Fig. 4A). Habitat heterogeneity also related positively to bird richness, but in a context-dependent way: the effect was most pronounced at intermediate levels of built-up cover and weakened to nearly flat in highly urbanised areas (Fig. 4B). In contrast, foliage-layer richness showed only a weak association with bird richness, with minor changes across its range and a slight decline that was most apparent in highly urbanised areas (Fig. 4C).

**Table 1.**
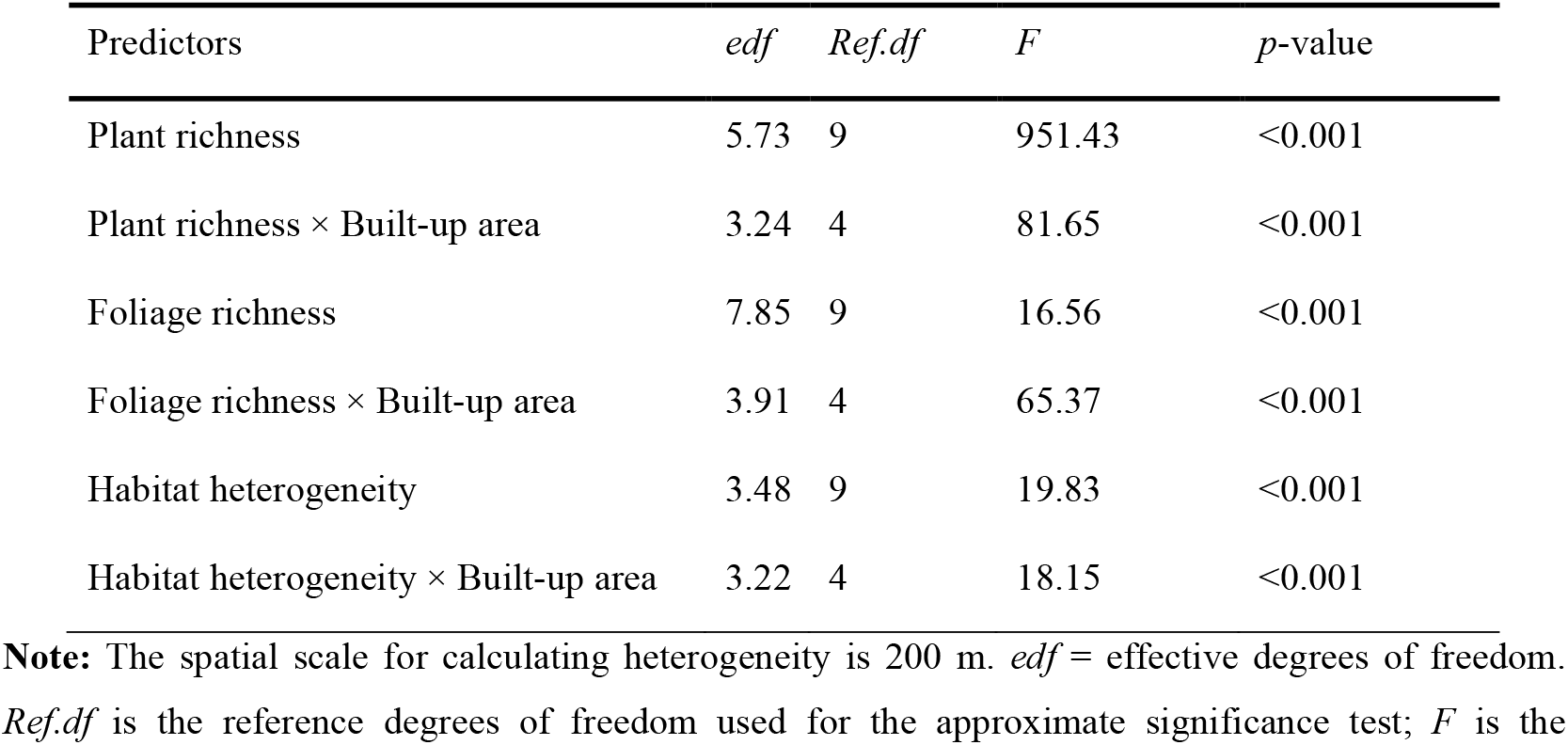
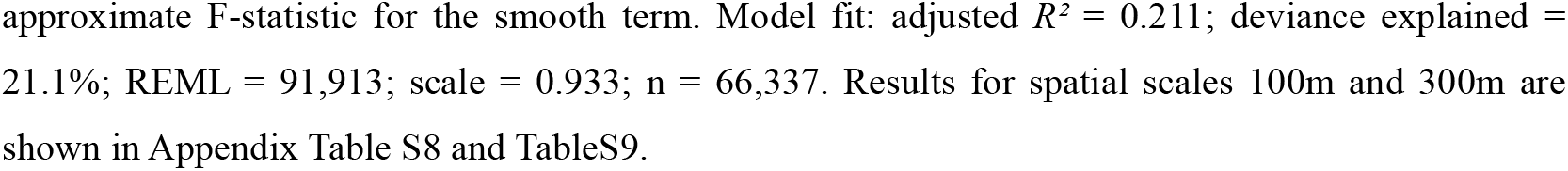
Generalized Additive Model (GAM) statistics for the effects of environmental heterogeneity on bird richness across urbanization intensity.

| Predictors | *edf* | *Ref.df* | *F* | *p*-value |
| --- | --- | --- | --- | --- |
| Plant richness | 5.73 | 9 | 951.43 | <0.001 |
| Plant richness × Built-up area | 3.24 | 4 | 81.65 | <0.001 |
| Foliage richness | 7.85 | 9 | 16.56 | <0.001 |
| Foliage richness × Built-up area | 3.91 | 4 | 65.37 | <0.001 |
| Habitat heterogeneity | 3.48 | 9 | 19.83 | <0.001 |
| Habitat heterogeneity × Built-up area | 3.22 | 4 | 18.15 | <0.001 |

**Note:** The spatial scale for calculating heterogeneity is 200 m. *edf* = effective degrees of freedom. *Ref.df* is the reference degrees of freedom used for the approximate significance test; *F* is the

**Figure 4.**
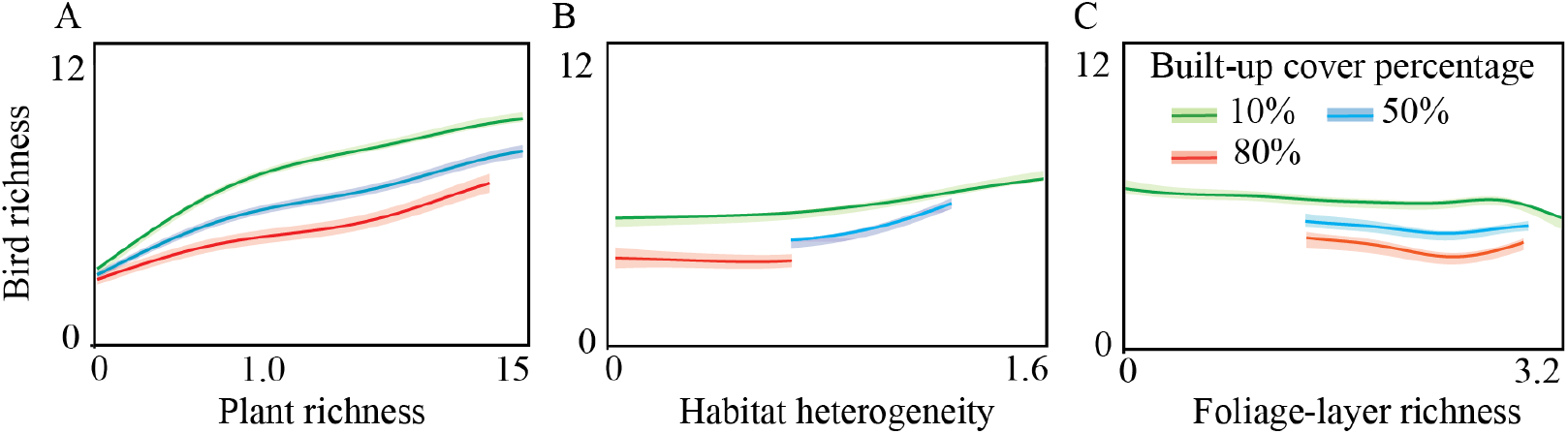
The effects of environmental heterogeneity on urban bird richness vary with urbanization intensity. (A–C) Conditional partial response curves from the same GAM showing how these relationships vary across the urbanisation gradient (built-up cover): (A) plant species richness, (B) habitat heterogeneity, and (C) foliage layer richness. Coloured lines indicate predictions at three representative built-up levels: 10% (green), 50% (blue), and 80% (orange); shaded ribbons show 95% confidence intervals. Bird richness was modelled as log (1 + richness), and plant species richness as ln (plant richness + 1), but axes are presented on the back-transformed (original) scales. The spatial scale for calculating heterogeneity is 200 m and results for spatial scale 100m and 300m are shown in Appendix Figure S3.

## 4. Discussion

Our study shows that three components of environmental heterogeneity, plant richness, habitat heterogeneity and foliage-layer richness, affect bird richness differently across the urbanization gradients. Plant species richness consistently supports higher bird richness across all urbanisation gradients, underscoring the importance of botanically diverse vegetation in urban landscapes. Habitat heterogeneity further enhances bird richness, with the strongest effects at intermediate built-up cover, whereas foliage-layer richness shows only limited effects. Urban bird richness declines with increasing built-up cover, underlining the pervasive negative effects of urbanization. However, this loss can be mitigated by context-dependent design of biotic environmental heterogeneity. We emphasize that conserving urban biodiversity requires providing biologically diverse and structurally heterogeneous habitat, with designs adapted to the local level of urbanisation.

Enhancing plant richness throughout the urbanization gradient can substantially promote urban bird richness. This association highlights the structuring role of plant diversity in urban ecosystems, as a richer plant species increases the diversity of food resources, nesting substrates, and shelter available to birds (Paker et al., 2014; Whelan & Maina, 2005). By expanding niche opportunities, higher plant richness can sustain the coexistence of both generalist and specialist bird species within urban landscapes (Kissling et al., 2008; Paker et al., 2014). The same level of plant richness may yield fewer bird species in highly built-up contexts than in greener landscapes, because green patches are typically smaller, more fragmented, and more isolated, and are embedded within a more hostile and disturbance-prone matrix (Allouche et al., 2012; Lamont & Pausas, 2024). Even so, increasing plant richness still remains a robust measure across different-level of urbanization intensities to meaningfully improve habitat quality and associated bird diversity.

Enhancing habitat heterogeneity supports urban bird diversity in greener areas, but its effectiveness declines in highly urbanized areas, emphasizing the importance of tailoring green-space design to urbanization intensity. The positive effects of habitat heterogeneity is supported by the habitat-heterogeneity hypothesis, which posits that greater environmental diversity expands niche space and resource opportunities, thereby promoting species coexistence (Anderle et al., 2023; Bazzaz, 1975; Tews et al., 2004). In urban systems, such heterogeneity was positively associated with bird richness in greener and moderately urbanized contexts, but this relationship weakened at very high built-up cover. This attenuation is consistent with the area–heterogeneity trade-off (Chiron et al., 2024). As urbanization increases, the total amount of vegetated habitat declines, and any additional heterogeneity is increasingly achieved by subdividing a limited green-space area into smaller and more isolated elements, reducing patch size and connectivity and constraining the number of individuals and species that can be supported (Allouche et al., 2012; Seiferling et al., 2014). The influence of habitat heterogeneity was strongest at intermediate levels of urbanization, where bird richness increased most sharply with heterogeneity. This pattern aligns with frameworks predicting unimodal relationships between environmental heterogeneity and biodiversity along human disturbance gradients, with diversity peaking at intermediate environmental complexity (Evans et al., 2007; Faeth et al., 2011; Seiferling et al., 2014; Sultana et al., 2021). In urban environments, such intermediate conditions likely allow the coexistence of both generalist and more specialized species: habitat diversity enhances resource complementarity and niche partitioning without yet crossing the fragmentation threshold that constrains richness (Callaghan et al., 2019). In planning terms, moderately urbanized areas appear to offer the greatest return on investment for enhancing habitat heterogeneity, with larger biodiversity gains than those achievable under highly built-up conditions.

Altering foliage-layer richness alone is unlikely to promote urban bird diversity, indicating that urban planning should prioritize factors such as plant species richness and habitat heterogeneity. A likely explanation is that additional foliage layers in urban green spaces often reflect intensively managed shrubs or ornamental plantings that increase vertical structure without delivering comparable gains for birds in food resources, nesting substrates, or microhabitat quality, particularly when woody species composition is simplified (Fontana et al., 2011). In some settings, greater layering may also indicate shrub and tree infilling that reduces open foraging space, potentially offsetting benefits for open-habitat guilds and producing a net neutral richness response. Consistent with this interpretation, bird responses to vertical structural metrics can be weak or neutral when structural complexity is decoupled from plant compositional diversity or coincides with habitat loss and fragmentation (Allouche et al., 2012; Breeuwer et al., 2009; Kissling et al., 2008). From an urban and landscape planning perspective, these results support prioritizing plant species richness and habitat heterogeneity, using vertical layering as a complementary design feature rather than a stand-alone biodiversity target.

For urban planning, our results suggest that citywide species-rich planting and targeted efforts to increase habitat heterogeneity in moderately built-up areas would greatly benefit bird richness. Below, we outline several actionable priorities for urban green space design. First, we recommend increasing the number of plant species during the design and management of urban green spaces. In practice, this can be achieved by encouraging spontaneous vegetation, planting diverse and functionally varied plant species (i.e., a combination of trees, shrubs, and herbaceous) and designing for species variation rather than uniform planting schemes. Second, we recommend urban planning to promote a well-connected network of diverse green features (e.g., diverse street trees, structurally varied gardens, and parks with small woodland patches). In practice, this means diversifying existing green spaces through management like retaining shrub patches, allowing seasonal “messier” areas, and maintaining hedgerows (Faeth et al., 2011; Jung et al., 2025).

## 5. Conclusion

This study shows that the effects of three types of environmental heterogeneity on bird species richness in urban landscapes vary along the level of urbanisation. Plant richness shows consistently positive effects the effect of increasing habitat heterogeneity is more pronounced in intermedium-level of urbanization, and. Plant species richness consistently supported higher bird richness, habitat heterogeneity was most beneficial at intermediate levels of urbanisation, whereas vegetation vertical structure heterogeneity shows limited influence. These findings indicate that sustaining urban bird diversity requires an urbanisation-intensity–dependent approach. Increasing plant species richness represents a broadly effective strategy across the urban gradient, while enhancing habitat heterogeneity is most effective in moderately urbanised landscapes. By demonstrating that different components of environmental heterogeneity have distinct effects along the urbanisation gradient, our study provides evidence-based guidance for designing and managing urban green spaces that support biodiversity across different levels of urban development.

## 6. Acknowledgements

J.H. acknowledges final project support from the China Scholarship Council (CSC, No. 202206180010). The contributions of J.M. and R.P.R. were supported by the Dutch Research Council (NWO) through the Merian Fund Cooperation China–The Netherlands (CAS) Green Cities 2019 (No. 482.19.704) and through COMBINED (NWA.1508.21.201). We thank the Leiden Institute for Environmental Science’s Impact Fund for the financial support provided during the preparation of this manuscript. This work was performed using the compute resources from the Academic Leiden Interdisciplinary Cluster Environment (ALICE) provided by Leiden University.

