## Appendix for "Designing urban green-space heterogeneity across urbanisation gradients to maintain avian richness"

**Table S1**: The classification of the Bird habitat type

| | Habitat type | Description of habitat type | Included LGN landcover classes and LGN number | | Definition of LGN landcover type |
|---|---|---|---|---|---|
| 1 | Herbaceous agriculture | All agricultural areas with low crops (herbaceous agriculture). | 1<br>2<br>3<br>4<br>5<br>6<br>10 | Agricultural grassland<br>Mais<br>Potato<br>Beetroot<br>Grain<br>Other agricultural crops<br>Flower bulbs | All agricultural classes are determined with the BRP and with Top10.<br>Regarding agricultural grassland, LGN does not include any grasslands in swamp areas, those fall under low shrubs in swamp. |
| 2 | Agricultural trees | All trees used for agricultural purposes. | 9<br>61<br>62 | Orchard<br>Tree nursery<br>Fruit tree plantation | All agricultural classes are determined with the BRP and with Top10.<br>Regarding agricultural grassland, LGN does not include any grasslands in swamp areas, those fall under low shrubs in swamp. |
| 3 | Build-up area | Buildings, greenhouses, bare ground, roads and railroads. | 18<br>19<br>24<br>25<br>26<br>27<br>8 | Building primary residential area<br>Building in secondary residential area<br>Bare ground in residential area<br>Roads and railroads<br>Buildings in suburban area<br>Other land use in suburbs<br>Greenhouses | • Buildings are taken from the Top10vlak, mask 'Stedelijk gebied'<br>• Infrastructure is taken from Top10<br>• Greenhouses are taken from Top10, at areas where the BRP does not show addresses<br>• Other land use in suburbs is based upon the Top10vlak 200 'overig grondgebruik'. Additionally, there must be no overlap with agricultural crops, swamp, marsh or peat. |
| 4 | Freshwater | All freshwater areas in the Netherlands. | 16 | Freshwater | This class is defined with the Top10 and BRP. |
| 5 | Saltwater | All saltwater areas in the Netherlands. | 17 | Saltwater | This class is defined with the Top10 and BRP. |
| 6 | Deciduous non-urban forest | Deciduous forest outside of urban, marsh, peat and swamp areas. | 11 | Deciduous forest | Classified with Top10 Nature mask and Top10 classes for deciduous and evergreen trees. They are not present in areas which are classified as marsh, peat or swamp. |
| 7 | Evergreen non-urban forest | Evergreen forest outside of urban, marsh, peat and swamp areas. | 12 | Evergreen forest | Classified with Top10 Nature mask and Top10 classes for deciduous and evergreen trees. They are not present in areas which are classified as marsh, peat or swamp. |
| 8 | Urban forest | All forests within residential areas, including both primary and secondary residential areas. | 20<br>22 | Forest in primary residential area<br>Forest in secondary residential area | Forests in residential area are regarded as all the forested areas that fall within the 'Stedelijk gebied' mask of Top10, AND are not included in swamp or peat areas.<br>This masks makes it possible to differentiate between primary and secondary residential areas. |
| 9 | Urban grassland | All grassland within residential areas, including both primary and secondary residential areas. | 23<br>28 | Grass in primary residential area<br>Grass in secondary residential area | Grasslands in residential area are regarded as all the forested areas that fall within the 'Stedelijk gebied' mask of Top10. This masks makes it possible to differentiate between primary and secondary residential areas. |
| 10 | Sand | All sandy areas within coastal | 31 | Open sand in coastal area | Appear only in coastal areas, taken as Top10 |

| | | | | | |
|---|---|---|---|---|---|
| | | areas and outside of coastal areas. | | | class 150: sand |
| | | | 35 | Driftsand | All areas that fall within the nature mask of Top10, area outside of coastal areas and are classified as sand in Top10. |
| 11 | Salt marsh | Salt marshes. | 30 | Salt marsh | They have a specific salt marsh mask for these areas |
| 12 | Dunes with low vegetation | Dunes with low vegetation. | 32 | Dunes with low vegetation | Low vegetation in coastal areas |
| 13 | Dunes with high vegetation | Dunes with high vegetation. | 33 | Dunes with high vegetation | High vegetation in coastal areas. |
| 14 | Heath | All heath landscape within and outside of coastal areas with less grass coverage than medium grassy heath areas. | 36 | Heath | Areas within Top10 swamp mask, which are not included in peat, natural grassland or marsh areas. |
| | | | 34 | Dune heath | All area present in the Top10 heath mask, and present in coastal areas. |
| 15 | Medium grassy heath | Heath area which is higher in grass composition than heath areas, and is separated from strong grassy heath by nature classification. | 37 | Medium grassy heath | Areas within Top10 swamp mask, which are not included in peat, natural grassland or marsh areas. |
| 16 | Strong grassy heath | Heath area which is higher in grass composition and has been separated through plant classification from medium grassy heath. | 38 | Strong grassy heath | Areas within Top10 swamp mask, which are not included in peat, natural grassland or marsh areas. |
| 17 | Low wetland shrubs | Low shrubs in wetland or swamp areas. | 322 | Low shrubs in swamp | Low vegetation in the Top10 swamp mask |
| | | | 41 | Other swamp vegetation | Low vegetation in the Top10 swamp mask. These areas were wrongfully classified as marsh areas, and have been reclassified by LGN as other swamp vegetation. |
| 18 | High wetland shrubs | High shrubs in wetland or swamp areas. This does not include trees. | 332 | High shrubs in swamp | High vegetation in the Top10 swamp mask. |
| 19 | Swamp forest | Forests within swamp areas. | 43 | Swamp forest | Trees in the Top10 swamp mask |
| 20 | Reed | Areas grown with reed. | 42 | Reed | Reeds in the Top10 swamp mask |
| 21 | Dry natural grassland | All dry grasslands which are outside of urban, agricultural and dune areas. These areas fall outside of wetland and peat areas. | 45 | Natural grassland | These are naturally governed grasslands. |
| | | | 46 | Coastal grassland | Grasslands which appear only in coastal areas of Top10. |
| | | | 47 | Other grassland | A category for all vegetation within the Top10 nature mask that is left after the other classifications. |
| 22 | Low dryland shrubs | Low shrubs outside of wetland, coastal and peat areas. | 323 | Other low shrubs | Low shrubs that are in the Top10 Nature mask, but do not fall under coastal, peat, swamp or marsh. |
| 23 | High dryland shrubs | High shrubs outside of wetland, coastal and peat areas. | 333 | Other high shrubs | High shrubs that are in the Top10 Nature mask, but do not fall under coastal, peat, swamp or marsh. |
| 24 | Peat moor | Peat moor areas without shrubs | 39 | Peat moor | Based upon the BRP classes and Top10 mask for peat areas. |
| 25 | Low peat moor shrubs | Peat moor areas with low shrub vegetation | 321 | Low peat moor shrubs | Based upon the BRP classes and Top10 mask for peat areas. |
| 26 | High peat moor shrubs | Peat moor areas with high shrub vegetation. | 331 | High peat moor shrubs | Based upon the BRP classes and Top10 mask for peat areas. |
| 27 | Peat moor forest | Forest in peat moor areas. | 40 | Peat moor forest | Based upon the BRP classes and Top10 mask for peat areas. |

**Table S2**: Bird species taxonomy and home-range radius reference dataset for setting analysis grid size

| **Dutch Name** | **Scientific name (NDFF)** | **Scientific name (e_Bird)** | **English name** | **Family2** | **Order2** | **Home range radius (km)** |
|---|---|---|---|---|---|---|
| Blauwe reiger | *Ardea cinerea* | *Ardea cinerea* | Gray Heron | Ardeidae | Pelecaniformes | |
| Zwartkop | *Sylvia atricapilla* | *Sylvia atricapilla* | Eurasian Blackcap | Sylviidae | Passeriformes | |
| Kolgans | *Anser albifrons* | *Anser albifrons* | Greater White-fronted Goose | Anatidae | Anseriformes | |
| Heggenmus | *Prunella modularis* | *Prunella modularis* | Dunnock | Prunellidae | Passeriformes | |
| Wilde eend | *Anas platyrhynchos* | *Anas platyrhyncho s* | Mallard | Anatidae | Anseriformes | |
| Grauwe klauwier | *Lanius collurio* | *Lanius collurio* | Red-backed Shrike | Laniidae | Passeriformes | 0.1256292 96 |
| Kleine karekiet | *Acrocephalus scirpaceus* | *Acrocephalus scirpaceus* | Eurasian Reed Warbler | Acrocephalidae | Passeriformes | |
| Turkse tortel | *Streptopelia decaocto* | *Streptopelia decaocto* | Eurasian Collared-Dove | Columbidae | Columbiformes | |
| Koolmees | *Parus major* | *Parus major* | Great Tit | Paridae | Passeriformes | |
| Pimpelmees | *Cyanistes caeruleus* | *Cyanistes caeruleus* | Eurasian Blue Tit | Paridae | Passeriformes | |
| Ekster | *Pica pica* | *Pica pica* | Eurasian Magpie | Corvidae | Passeriformes | |
| Houtduif | *Columba palumbus* | *Columba palumbus* | Common Wood-Pigeon | Columbidae | Columbiformes | 1.5937377 45 |
| Groenling | *Chloris chloris* | *Chloris chloris* | European Greenfinch | Fringillidae | Passeriformes | |
| Merel | *Turdus merula* | *Turdus merula* | Common Blackbird | Turdidae | Passeriformes | |
| Knobbelzwaan | *Cygnus olor* | *Cygnus olor* | Mute Swan | Anatidae | Anseriformes | |
| Kruisbek | *Loxia curvirostra* | *Loxia curvirostra* | Common Crossbill | Fringillidae | Passeriformes | |
| Roek | *Corvus frugilegus* | *Corvus frugilegus* | Rook | Corvidae | Passeriformes | |
| Putter | *Carduelis carduelis* | *Carduelis carduelis* | European Goldfinch | Fringillidae | Passeriformes | |
| Kraanvogel | *Grus grus* | *Grus grus* | Common Crane | Gruidae | Gruiformes | |
| Havik | *Accipiter gentilis* | *Accipiter gentilis* | Northern Goshawk | Accipitridae | Accipitriformes | 6.3245553 2 |
| Tjiftjaf | *Phylloscopus collybita* | *Phylloscopus collybita* | Common Chiffchaff | Phylloscopidae | Passeriformes | |
| Koekoek | *Cuculus canorus* | *Cuculus canorus* | Common Cuckoo | Cuculidae | Cuculiformes | 6.2016126 93 |
| Huismus | *Passer domesticus* | *Passer domesticus* | House Sparrow | Passeridae | Passeriformes | |
| Groene specht | *Picus viridis* | *Picus viridis* | European Green Woodpecker | Picidae | Piciformes | 1.3601470 51 |
| Gierzwaluw | *Apus apus* | *Apus apus* | Common Swift | Apodidae | Caprimulgiform es | |
| Raaf | *Corvus corax* | *Corvus corax* | Common Raven | Corvidae | Passeriformes | 5.2915026 22 |

| Dutch Name | Scientific name (NDFF) | Scientific name (e_Bird) | English name | Family2 | Order2 | Home range radius (km) |
|---|---|---|---|---|---|---|
| Boomvalk | *Falco subbuteo* | *Falco subbuteo* | Eurasian Hobby | Falconidae | Falconiformes | |
| Sperwer | *Accipiter nisus* | *Accipiter nisus* | Eurasian Sparrowhawk | Accipitridae | Accipitriformes | 2.664582519 |
| Torenvalk | *Falco tinnunculus* | *Falco tinnunculus* | Common Kestrel | Falconidae | Falconiformes | 1.732050808 |
| Boerenzwaluw | *Hirundo rustica* | *Hirundo rustica* | Barn Swallow | Hirundinidae | Passeriformes | |
| Zwarte kraai | *Corvus corone* | *Corvus corone* | Carrion Crow | Corvidae | Passeriformes | |
| Kleine mantelmeeuw | *Larus fuscus* | *Larus fuscus* | Lesser Black-backed Gull | Laridae | Charadriiformes | |
| Grutto | *Limosa limosa* | *Limosa limosa* | Black-tailed Godwit | Scolopacidae | Charadriiformes | |
| Kluut | *Recurvirostra avosetta* | *Recurvirostra avosetta* | Pied Avocet | Recurvirostridae | Charadriiformes | |
| Bonte strandloper | *Calidris alpina* | *Calidris alpina* | Dunlin | Scolopacidae | Charadriiformes | |
| Goudvink | *Pyrrhula pyrrhula* | *Pyrrhula pyrrhula* | Eurasian Bullfinch | Fringillidae | Passeriformes | |
| Kauw | *Corvus monedula* | *Corvus monedula* | Eurasian Jackdaw | Corvidae | Passeriformes | |
| Boomkruiper | *Certhia brachydactyla* | *Certhia brachydactyla* | Short-toed Treecreeper | Certhiidae | Passeriformes | |
| Grote bonte specht | *Dendrocopos major* | *Dendrocopos major* | Great Spotted Woodpecker | Picidae | Piciformes | |
| Bosrietzanger | *Acrocephalus palustris* | *Acrocephalus palustris* | Marsh Warbler | Acrocephalidae | Passeriformes | |
| Zeearend | *Haliaeetus albicilla* | *Haliaeetus albicilla* | White-tailed Eagle | Accipitridae | Accipitriformes | |
| Huiszwaluw | *Delichon urbicum* | *Delichon urbicum* | Common House-Martin | Hirundinidae | Passeriformes | |
| Wilde zwaan | *Cygnus cygnus* | *Cygnus cygnus* | Whooper Swan | Anatidae | Anseriformes | |
| Buizerd | *Buteo buteo* | *Buteo buteo* | Common Buzzard | Accipitridae | Accipitriformes | 7.088018059 |
| Nijlgans | *Alopochen aegyptiaca* | *Alopochen aegyptiaca* | Egyptian Goose | Anatidae | Anseriformes | |
| Bruine kiekendief | *Circus aeruginosus* | *Circus aeruginosus* | Eurasian Marsh Harrier | Accipitridae | Accipitriformes | |
| Watersnip | *Gallinago gallinago* | *Gallinago gallinago* | Common Snipe | Scolopacidae | Charadriiformes | |
| Spreeuw | *Sturnus vulgaris* | *Sturnus vulgaris* | European Starling | Sturnidae | Passeriformes | |
| Brilduiker | *Bucephala clangula* | *Bucephala clangula* | Common Goldeneye | Anatidae | Anseriformes | |
| Middelste zaagbek | *Mergus serrator* | *Mergus serrator* | Red-breasted Merganser | Anatidae | Anseriformes | |
| IJseend | *Clangula hyemalis* | *Clangula hyemalis* | Long-tailed Duck | Anatidae | Anseriformes | |
| Grote stern | *Sterna sandvicensis* | *Thalasseus sandvicensis* | Sandwich Tern | Laridae | Charadriiformes | |
| IJsvogel | *Alcedo atthis* | *Alcedo atthis* | Common Kingfisher | Alcedinidae | Coraciiformes | |

| Dutch Name | Scientific name (NDFF) | Scientific name (e_Bird) | English name | Family2 | Order2 | Home range radius (km) |
|---|---|---|---|---|---|---|
| Dodaars | *Tachybaptus ruficollis* | *Tachybaptus ruficollis* | Little Grebe | Podicipedidae | Podicipediformes | |
| Meerkoet | *Fulica atra* | *Fulica atra* | Eurasian Coot | Rallidae | Gruiformes | |
| Slechtvalk | *Falco peregrinus* | *Falco peregrinus* | Peregrine Falcon | Falconidae | Falconiformes | 12.4040316 |
| Boomklever | *Sitta europaea* | *Sitta europaea* | Eurasian Nuthatch | Sittidae | Passeriformes | 0.144913767 |
| Grote lijster | *Turdus viscivorus* | *Turdus viscivorus* | Mistle Thrush | Turdidae | Passeriformes | |
| Staartmees | *Aegithalos caudatus* | *Aegithalos caudatus* | Long-tailed Tit | Aegithalidae | Passeriformes | 0.204939015 |
| Grote zaagbek | *Mergus merganser* | *Mergus merganser* | Common Merganser | Anatidae | Anseriformes | |
| Nonnetje | *Mergellus albellus* | *Mergellus albellus* | Smew | Anatidae | Anseriformes | |
| Oeverzwaluw | *Riparia riparia* | *Riparia riparia* | Sand Martin | Hirundinidae | Passeriformes | |
| Gekraagde roodstaart | *Phoenicurus phoenicurus* | *Phoenicurus phoenicurus* | Common Redstart | Muscicapidae | Passeriformes | 0.067082039 |
| Tapuit | *Oenanthe oenanthe* | *Oenanthe oenanthe* | Northern Wheatear | Muscicapidae | Passeriformes | 0.124008185 |
| Krooneend | *Netta rufina* | *Netta rufina* | Red-crested Pochard | Anatidae | Anseriformes | |
| Rietgors | *Emberiza schoeniclus* | *Emberiza schoeniclus* | Common Reed Bunting | Emberizidae | Passeriformes | |
| Witte kwikstaart | *Motacilla alba* | *Motacilla alba* | White Wagtail | Motacillidae | Passeriformes | 0.886002257 |
| Rode wouw | *Milvus milvus* | *Milvus milvus* | Red Kite | Accipitridae | Accipitriformes | 4.430011287 |
| Grauwe gans | *Anser anser* | *Anser anser* | Greylag Goose | Anatidae | Anseriformes | |
| Rosse stekelstaart | *Oxyura jamaicensis* | *Oxyura jamaicensis* | Ruddy Duck | Anatidae | Anseriformes | |
| Drieteenmeeuw | *Rissa tridactyla* | *Rissa tridactyla* | Black-legged Kittiwake | Laridae | Charadriiformes | |
| Witgat | *Tringa ochropus* | *Tringa ochropus* | Green Sandpiper | Scolopacidae | Charadriiformes | |
| Kokmeeuw | *Chroicocephalus ridibundus* | *Chroicocephalus ridibundus* | Black-headed Gull | Laridae | Charadriiformes | |
| Winterkoning | *Troglodytes troglodytes* | *Troglodytes troglodytes* | Eurasian Wren | Troglodytidae | Passeriformes | 0.100583945 |
| Kievit | *Vanellus vanellus* | *Vanellus vanellus* | Northern Lapwing | Charadriidae | Charadriiformes | |
| Bergeend | *Tadorna tadorna* | *Tadorna tadorna* | Common Shelduck | Anatidae | Anseriformes | |
| Brandgans | *Branta leucopsis* | *Branta leucopsis* | Barnacle Goose | Anatidae | Anseriformes | |
| Tureluur | *Tringa totanus* | *Tringa totanus* | Common Redshank | Scolopacidae | Charadriiformes | |
| Stormmeeuw | *Larus canus* | *Larus canus* | Common Gull | Laridae | Charadriiformes | |
| Zilvermeeuw | *Larus argentatus* | *Larus argentatus* | European Herring Gull | Laridae | Charadriiformes | |
| Veldleeuwerik | *Alauda arvensis* | *Alauda arvensis* | Eurasian Skylark | Alaudidae | Passeriformes | |
| Smient | *Anas penelope* | *Mareca* | Eurasian | Anatidae | Anseriformes | |

| Dutch Name | Scientific name (NDFF) | Scientific name (e_Bird) | English name | Family2 | Order2 | Home range radius (km) |
|---|---|---|---|---|---|---|
| | | *penelope* | Wigeon | | | |
| Wintertaling | *Anas crecca* | *Anas crecca* | Green-winged Teal | Anatidae | Anseriformes | |
| Ringmus | *Passer montanus* | *Passer montanus* | Eurasian Tree Sparrow | Passeridae | Passeriformes | |
| Holenduif | *Columba oenas* | *Columba oenas* | Stock Dove | Columbidae | Columbiformes | |
| Keep | *Fringilla montifringilla* | *Fringilla montifringilla* | Brambling | Fringillidae | Passeriformes | |
| Tafeleend | *Aythya ferina* | *Aythya ferina* | Common Pochard | Anatidae | Anseriformes | |
| Rotgans | *Branta bernicla* | *Branta bernicla* | Brant | Anatidae | Anseriformes | |
| Aalscholver | *Phalacrocorax carbo* | *Phalacrocorax carbo* | Great Cormorant | Phalacrocoracidae | Suliformes | |
| Wulp | *Numenius arquata* | *Numenius arquata* | Eurasian Curlew | Scolopacidae | Charadriiformes | |
| Goudplevier | *Pluvialis apricaria* | *Pluvialis apricaria* | European Golden-Plover | Charadriidae | Charadriiformes | |
| Grote gele kwikstaart | *Motacilla cinerea* | *Motacilla cinerea* | Grey Wagtail | Motacillidae | Passeriformes | |
| Zanglijster | *Turdus philomelos* | *Turdus philomelos* | Song Thrush | Turdidae | Passeriformes | |
| Fuut | *Podiceps cristatus* | *Podiceps cristatus* | Great Crested Grebe | Podicipedidae | Podicipediformes | |
| Kuifeend | *Aythya fuligula* | *Aythya fuligula* | Tufted Duck | Anatidae | Anseriformes | |
| Pijlstaart | *Anas acuta* | *Anas acuta* | Northern Pintail | Anatidae | Anseriformes | |
| Krakeend | *Anas strepera* | *Mareca strepera* | Gadwall | Anatidae | Anseriformes | |
| Baardman | *Panurus biarmicus* | *Panurus biarmicus* | Bearded Reedling | Panuridae | Passeriformes | |
| Slobeend | *Anas clypeata* | *Spatula clypeata* | Northern Shoveler | Anatidae | Anseriformes | |
| Scholekster | *Haematopus ostralegus* | *Haematopus ostralegus* | Eurasian Oystercatcher | Haematopodidae | Charadriiformes | 1.568438714 |
| Geoorde fuut | *Podiceps nigricollis* | *Podiceps nigricollis* | Black-necked Grebe | Podicipedidae | Podicipediformes | |
| Visdief | *Sterna hirundo* | *Sterna hirundo* | Common Tern | Laridae | Charadriiformes | |
| Rietzanger | *Acrocephalus schoenobaenus* | *Acrocephalus schoenobaenus* | Sedge Warbler | Acrocephalidae | Passeriformes | |
| Snor | *Locustella luscinioides* | *Locustella luscinioides* | Savi's Warbler | Locustellidae | Passeriformes | |
| Blauwe kiekendief | *Circus cyaneus* | *Circus cyaneus* | Northern Harrier | Accipitridae | Accipitriformes | 1.587824786 |
| Blauwborst | *Luscinia svecica* | *Luscinia svecica* | Bluethroat | Muscicapidae | Passeriformes | |
| Roodborst | *Erithacus rubecula* | *Erithacus rubecula* | European Robin | Muscicapidae | Passeriformes | |
| Waterpieper | *Anthus spinoletta* | *Anthus spinoletta* | Water Pipit | Motacillidae | Passeriformes | |
| Waterhoen | *Gallinula chloropus* | *Gallinula chloropus* | Common Moorhen | Rallidae | Gruiformes | |

| Dutch Name | Scientific name (NDFF) | Scientific name (e_Bird) | English name | Family2 | Order2 | Home range radius (km) |
|---|---|---|---|---|---|---|
| Graspieper | *Anthus pratensis* | *Anthus pratensis* | Meadow Pipit | Motacillidae | Passeriformes | |
| Rosse grutto | *Limosa lapponica* | *Limosa lapponica* | Bar-tailed Godwit | Scolopacidae | Charadriiformes | |
| Grote mantelmeeuw | *Larus marinus* | *Larus marinus* | Great Black-backed Gull | Laridae | Charadriiformes | |
| Dwergmeeuw | *Hydrocoloeus minutus* | *Hydrocoloeus minutus* | Little Gull | Laridae | Charadriiformes | |
| Bontbekplevier | *Charadrius hiaticula* | *Charadrius hiaticula* | Common Ringed Plover | Charadriidae | Charadriiformes | |
| Kramsvogel | *Turdus pilaris* | *Turdus pilaris* | Fieldfare | Turdidae | Passeriformes | |
| Boomleeuwerik | *Lullula arborea* | *Lullula arborea* | Woodlark | Alaudidae | Passeriformes | |
| Roodkeelduiker | *Gavia stellata* | *Gavia stellata* | Red-throated Loon | Gaviidae | Gaviiformes | |
| Kuifduiker | *Podiceps auritus* | *Podiceps auritus* | Horned Grebe | Podicipedidae | Podicipediformes | |
| Jan-van-gent | *Morus bassanus* | *Morus bassanus* | Northern Gannet | Sulidae | Suliformes | |
| Grasmus | *Sylvia communis* | *Curruca communis* | Common Whitethroat | Sylviidae | Passeriformes | |
| Groenpootruiter | *Tringa nebularia* | *Tringa nebularia* | Common Greenshank | Scolopacidae | Charadriiformes | |
| Buffelkopeend | *Bucephala albeola* | *Bucephala albeola* | Bufflehead | Anatidae | Anseriformes | |
| Oeverloper | *Actitis hypoleucos* | *Actitis hypoleucos* | Common Sandpiper | Scolopacidae | Charadriiformes | |
| Paapje | *Saxicola rubetra* | *Saxicola rubetra* | Whinchat | Muscicapidae | Passeriformes | 0.085440037 |
| Paarse strandloper | *Calidris maritima* | *Calidris maritima* | Purple Sandpiper | Scolopacidae | Charadriiformes | |
| Zilverplevier | *Pluvialis squatarola* | *Pluvialis squatarola* | Grey Plover | Charadriidae | Charadriiformes | |
| Kleine zwaan | *Cygnus bewickii* | *Cygnus columbianus* | Tundra Swan | Anatidae | Anseriformes | |
| Koperwiek | *Turdus iliacus* | *Turdus iliacus* | Redwing | Turdidae | Passeriformes | |
| Vuurgoudhaan | *Regulus ignicapilla* | *Regulus ignicapilla* | Common Firecrest | Regulidae | Passeriformes | |
| Gaai | *Garrulus glandarius* | *Garrulus glandarius* | Eurasian Jay | Corvidae | Passeriformes | |
| Vink | *Fringilla coelebs* | *Fringilla coelebs* | Common Chaffinch | Fringillidae | Passeriformes | 0.204939015 |
| Eider | *Somateria mollissima* | *Somateria mollissima* | Common Eider | Anatidae | Anseriformes | |
| Roodhalsfuut | *Podiceps grisegena* | *Podiceps grisegena* | Red-necked Grebe | Podicipedidae | Podicipediformes | |
| Buidelmees | *Remiz pendulinus* | *Remiz pendulinus* | Eurasian Penduline-Tit | Remizidae | Passeriformes | |
| Appelvink | *Coccothraustes coccothraustes* | *Coccothraustes coccothraustes* | European Hawfinch | Fringillidae | Passeriformes | |
| Lepelaar | *Platalea leucorodia* | *Platalea leucorodia* | Eurasian Spoonbill | Threskiornithidae | Pelecaniformes | |
| Regenwulp | *Numenius* | *Numenius* | Whimbrel | Scolopacidae | Charadriiformes | |

| Dutch Name | Scientific name (NDFF) | Scientific name (e_Bird) | English name | Family2 | Order2 | Home range radius (km) |
|---|---|---|---|---|---|---|
| | *phaeopus* | *phaeopus* | | | | |
| Tuinfluiter | *Sylvia borin* | *Sylvia borin* | Garden Warbler | Sylviidae | Passeriformes | |
| Klapekster | *Lanius excubitor* | *Lanius excubitor* | Great Gray Shrike | Laniidae | Passeriformes | |
| Fitis | *Phylloscopus trochilus* | *Phylloscopus trochilus* | Willow Warbler | Phylloscopidae | Passeriformes | |
| Ooievaar | *Ciconia ciconia* | *Ciconia ciconia* | White Stork | Ciconiidae | Ciconiiformes | |
| Steltkluut | *Himantopus himantopus* | *Himantopus himantopus* | Black-winged Stilt | Recurvirostridae | Charadriiformes | |
| Zwarte ruiter | *Tringa erythropus* | *Tringa erythropus* | Spotted Redshank | Scolopacidae | Charadriiformes | |
| Waterral | *Rallus aquaticus* | *Rallus aquaticus* | Water Rail | Rallidae | Gruiformes | |
| Smelleken | *Falco columbarius* | *Falco columbarius* | Merlin | Falconidae | Falconiformes | |
| Matkop | *Poecile montanus* | *Poecile montanus* | Willow Tit | Paridae | Passeriformes | |
| Roodborsttapuit | *Saxicola rubicola* | *Saxicola rubicola* | European Stonechat | Muscicapidae | Passeriformes | |
| Ruigpootbuizerd | *Buteo lagopus* | *Buteo lagopus* | Rough-legged Hawk | Accipitridae | Accipitriformes | |
| Gele kwikstaart | *Motacilla flava* | *Motacilla flava* | Western Yellow Wagtail | Motacillidae | Passeriformes | 0.100583945 |
| Goudhaan | *Regulus regulus* | *Regulus regulus* | Goldcrest | Regulidae | Passeriformes | 0.14106736 |
| Boompieper | *Anthus trivialis* | *Anthus trivialis* | Tree Pipit | Motacillidae | Passeriformes | |
| Geelgors | *Emberiza citrinella* | *Emberiza citrinella* | Yellowhammer | Emberizidae | Passeriformes | |
| Roodhalsgans | *Branta ruficollis* | *Branta ruficollis* | Red-breasted Goose | Anatidae | Anseriformes | |
| Zwarte Zee-eend | *Melanitta nigra* | *Melanitta nigra* | Common Scoter | Anatidae | Anseriformes | |
| Casarca | *Tadorna ferruginea* | *Tadorna ferruginea* | Ruddy Shelduck | Anatidae | Anseriformes | |
| Fazant | *Phasianus colchicus* | *Phasianus colchicus* | Ring-necked Pheasant | Phasianidae | Galliformes | |
| Zwarte roodstaart | *Phoenicurus ochruros* | *Phoenicurus ochruros* | Black Redstart | Muscicapidae | Passeriformes | |
| Glanskop | *Poecile palustris* | *Poecile palustris* | Marsh Tit | Paridae | Passeriformes | |
| Wespendief | *Pernis apivorus* | *Pernis apivorus* | European Honey-buzzard | Accipitridae | Accipitriformes | |
| Purperreiger | *Ardea purpurea* | *Ardea purpurea* | Purple Heron | Ardeidae | Pelecaniformes | |
| Zomertaling | *Anas querquedula* | *Spatula querquedula* | Garganey | Anatidae | Anseriformes | |
| Kleine plevier | *Charadrius dubius* | *Charadrius dubius* | Little Ringed Plover | Charadriidae | Charadriiformes | |
| Grote pieper | *Anthus richardi* | *Anthus richardi* | Richard’s Pipit | Motacillidae | Passeriformes | |
| Visarend | *Pandion haliaetus* | *Pandion haliaetus* | Osprey | Pandionidae | Accipitriformes | |

| Dutch Name | Scientific name (NDFF) | Scientific name (e_Bird) | English name | Family2 | Order2 | Home range radius (km) |
|---|---|---|---|---|---|---|
| Grauwe franjepoot | *Phalaropus lobatus* | *Phalaropus lobatus* | Red-necked Phalarope | Scolopacidae | Charadriiformes | |
| Zwarte stern | *Chlidonias niger* | *Chlidonias niger* | Black Tern | Laridae | Charadriiformes | |
| Kleine zilverreiger | *Egretta garzetta* | *Egretta garzetta* | Little Egret | Ardeidae | Pelecaniformes | |
| Roerdomp | *Botaurus stellaris* | *Botaurus stellaris* | Great Bittern | Ardeidae | Pelecaniformes | 0.439317653 |
| Kleine strandloper | *Calidris minuta* | *Calidris minuta* | Little Stint | Scolopacidae | Charadriiformes | |
| Steenloper | *Arenaria interpres* | *Arenaria interpres* | Ruddy Turnstone | Scolopacidae | Charadriiformes | |
| Kwartel | *Coturnix coturnix* | *Coturnix coturnix* | Common Quail | Phasianidae | Galliformes | |
| Porseleinhoen | *Porzana porzana* | *Porzana porzana* | Spotted Crake | Rallidae | Gruiformes | |
| Zwarthalszwaan | *Cygnus melancoryphus* | *Cygnus melancoryphus* | Black-necked Swan | Anatidae | Anseriformes | |
| Cetti's Zanger | *Cettia cetti* | *Cettia cetti* | Cetti’s Warbler | Scotocercidae | Passeriformes | |
| Kanoet | *Calidris canutus* | *Calidris canutus* | Red Knot | Scolopacidae | Charadriiformes | |
| Steppekiekendief | *Circus macrourus* | *Circus macrourus* | Pallid Harrier | Accipitridae | Accipitriformes | |
| Kleine rietgans | *Anser brachyrhynchus* | *Anser brachyrhynchus* | Pink-footed Goose | Anatidae | Anseriformes | |
| Sprinkhaanzanger | *Locustella naevia* | *Locustella naevia* | Common Grasshopper-Warbler | Locustellidae | Passeriformes | |
| Witvleugelstern | *Chlidonias leucopterus* | *Chlidonias leucopterus* | White-winged Tern | Laridae | Charadriiformes | |
| Flamingo | *Phoenicopterus roseus* | *Phoenicopterus roseus* | Greater Flamingo | Phoenicopteridae | Phoenicopteriformes | |
| Koereiger | *Bubulcus ibis* | *Bubulcus ibis* | Cattle Egret | Ardeidae | Pelecaniformes | |
| Ransuil | *Asio otus* | *Asio otus* | Long-eared Owl | Strigidae | Strigiformes | 4.429446918 |
| Indische gans | *Anser indicus* | *Anser indicus* | Bar-headed Goose | Anatidae | Anseriformes | |
| Kuifmees | *Lophophanes cristatus* | *Lophophanes cristatus* | European Crested Tit | Paridae | Passeriformes | |
| Nachtegaal | *Luscinia megarhynchos* | *Luscinia megarhynchos* | Common Nightingale | Muscicapidae | Passeriformes | |
| Braamsluiper | *Sylvia curruca* | *Curruca curruca* | Lesser Whitethroat | Sylviidae | Passeriformes | |
| Dwergstern | *Sternula albifrons* | *Sternula albifrons* | Little Tern | Laridae | Charadriiformes | |
| Wielewaal | *Oriolus oriolus* | *Oriolus oriolus* | Eurasian Golden Oriole | Oriolidae | Passeriformes | |
| Beflijster | *Turdus torquatus* | *Turdus torquatus* | Ring Ouzel | Turdidae | Passeriformes | |
| Noordse stern | *Sterna paradisaea* | *Sterna paradisaea* | Arctic Tern | Laridae | Charadriiformes | |
| Chileense flamingo | *Phoenicopterus chilensis* | *Phoenicopterus chilensis* | Chilean Flamingo | Phoenicopteridae | Phoenicopteriformes | |
| Zomertortel | *Streptopelia turtur* | *Streptopelia turtur* | European Turtle-Dove | Columbidae | Columbiformes | 7.974020316 |

| Dutch Name | Scientific name (NDFF) | Scientific name (e_Bird) | English name | Family2 | Order2 | Home range radius (km) |
|---|---|---|---|---|---|---|
| Bonapartes strandloper | *Calidris fuscicollis* | *Calidris fuscicollis* | White-rumped Sandpiper | Scolopacidae | Charadriiformes | |
| Patrijs | *Perdix perdix* | *Perdix perdix* | Grey Partridge | Phasianidae | Galliformes | 0.248997992 |
| Zwarte ibis | *Plegadis falcinellus* | *Plegadis falcinellus* | Glossy Ibis | Threskiornithidae | Pelecaniformes | |
| Zwarte rotgans | *Branta nigricans* | *Branta bernicla* | Brant | Anatidae | Anseriformes | |
| Bonte vliegenvanger | *Ficedula hypoleuca* | *Ficedula hypoleuca* | European Pied Flycatcher | Muscicapidae | Passeriformes | |
| Spotvogel | *Hippolais icterina* | *Hippolais icterina* | Icterine Warbler | Acrocephalidae | Passeriformes | |
| Drieteenstrandloper | *Calidris alba* | *Calidris alba* | Sanderling | Scolopacidae | Charadriiformes | |
| Zwarte ooievaar | *Ciconia nigra* | *Ciconia nigra* | Black Stork | Ciconiidae | Ciconiiformes | |
| Halsbandparkiet | *Psittacula krameri* | *Psittacula krameri* | Rose-ringed Parakeet | Psittaculidae | Psittaciformes | |
| Rosse franjepoot | *Phalaropus fulicarius* | *Phalaropus fulicarius* | Red Phalarope | Scolopacidae | Charadriiformes | |
| Oeverpieper | *Anthus petrosus* | *Anthus petrosus* | Rock Pipit | Motacillidae | Passeriformes | |
| Parelduiker | *Gavia arctica* | *Gavia arctica* | Black-throated Loon | Gaviidae | Gaviiformes | |
| Zeekoet | *Uria aalge* | *Uria aalge* | Common Murre | Alcidae | Charadriiformes | |
| Bladkoning | *Phylloscopus inornatus* | *Phylloscopus inornatus* | Yellow-browed Warbler | Phylloscopidae | Passeriformes | |
| Zwarte zwaan | *Cygnus atratus* | *Cygnus atratus* | Black Swan | Anatidae | Anseriformes | |
| Sneeuwgors | *Plectrophenax nivalis* | *Plectrophenax nivalis* | Snow Bunting | Calcariidae | Passeriformes | |
| IJslandse grutto | *Limosa limosa islandica* | *Limosa limosa* | Black-tailed Godwit | Scolopacidae | Charadriiformes | |
| Kleine jager | *Stercorarius parasiticus* | *Stercorarius parasiticus* | Parasitic Jaeger | Stercorariidae | Charadriiformes | |
| Zwartkopmeeuw | *Larus melanocephalus* | *Ichthyaetus melanocephalus* | Mediterranean Gull | Laridae | Charadriiformes | |
| Kleine geelpootruiter | *Tringa flavipes* | *Tringa flavipes* | Lesser Yellowlegs | Scolopacidae | Charadriiformes | |
| Europese kanarie | *Serinus serinus* | *Serinus serinus* | European Serin | Fringillidae | Passeriformes | 0.1 |
| Grauwe vliegenvanger | *Muscicapa striata* | *Muscicapa striata* | Spotted Flycatcher | Muscicapidae | Passeriformes | 0.1 |
| Topper | *Aythya marila* | *Aythya marila* | Greater Scaup | Anatidae | Anseriformes | |
| Roze pelikaan | *Pelecanus onocrotalus* | *Pelecanus onocrotalus* | Great White Pelican | Pelecanidae | Pelecaniformes | |
| Pontische meeuw | *Larus cachinnans* | *Larus cachinnans* | Caspian Gull | Laridae | Charadriiformes | |
| Bosuil | *Strix aluco* | *Strix aluco* | Tawny Owl | Strigidae | Strigiformes | 0.597438005 |
| Grote Zee-eend | *Melanitta fusca* | *Melanitta fusca* | Velvet Scoter | Anatidae | Anseriformes | |
| Kerkuil | *Tyto alba* | *Tyto alba* | Barn Owl | Tytonidae | Strigiformes | 1.224744871 |
| Oehoe | *Bubo bubo* | *Bubo bubo* | Eurasian | Strigidae | Strigiformes | 4 |

| Dutch Name | Scientific name (NDFF) | Scientific name (e_Bird) | English name | Family2 | Order2 | Home range radius (km) |
|---|---|---|---|---|---|---|
| | | | Eagle-Owl | | | |
| Zwarte mees | *Periparus ater* | *Periparus ater* | Coal Tit | Paridae | Passeriformes | |
| Witkopstaartmees | *Aegithalos caudatus caudatus* | *Aegithalos caudatus* | Long-tailed Tit | Aegithalidae | Passeriformes | 0.204939015 |
| Stadsduif | *Columba livia f. domestica* | *Columba livia* | Rock Pigeon | Columbidae | Columbiformes | |
| Houtsnip | *Scolopax rusticola* | *Scolopax rusticola* | Eurasian Woodcock | Scolopacidae | Charadriiformes | |
| Zwarte zeekoet | *Cepphus grylle* | *Cepphus grylle* | Black Guillemot | Alcidae | Charadriiformes | |
| Grote alexanderparkiet | *Psittacula eupatria* | *Psittacula eupatria* | Alexandrine Parakeet | Psittaculidae | Psittaciformes | |
| Rouwkwikstaart | *Motacilla yarrellii* | *Motacilla alba* | White Wagtail | Motacillidae | Passeriformes | 0.886002257 |
| Muskuseend | *Cairina moschata* | *Cairina moschata* | Muscovy Duck | Anatidae | Anseriformes | |
| Geelpootmeeuw | *Larus michahellis* | *Larus michahellis* | Yellow-legged Gull | Laridae | Charadriiformes | |
| Zwarte specht | *Dryocopus martius* | *Dryocopus martius* | Black Woodpecker | Picidae | Piciformes | 1.870828693 |
| Morinelplevier | *Charadrius morinellus* | *Charadrius morinellus* | Eurasian Dotterel | Charadriidae | Charadriiformes | |
| Grote jager | *Stercorarius skua* | *Stercorarius skua* | Great Skua | Phalacrocoracidae | Suliformes | |
| Kuifaalscholver | *Phalacrocorax aristotelis* | *Gulosus aristotelis* | European Shag | Phalacrocoracidae | Suliformes | |
| IJsduiker | *Gavia immer* | *Gavia immer* | Common Loon | Gaviidae | Gaviiformes | |
| Mandarijneend | *Aix galericulata* | *Aix galericulata* | Mandarin Duck | Anatidae | Anseriformes | |
| Engelse kwikstaart | *Motacilla flavissima* | *Motacilla flava* | Western Yellow Wagtail | Motacillidae | Passeriformes | 0.100583945 |
| Velduil | *Asio flammeus* | *Asio flammeus* | Short-eared Owl | Strigidae | Strigiformes | |
| Amerikaanse wintertaling | *Anas carolinensis* | *Anas crecca* | Green-winged Teal | Anatidae | Anseriformes | |
| Carolina-eend | *Aix sponsa* | *Aix sponsa* | Wood Duck | Anatidae | Anseriformes | |
| IJsgors | *Calcarius lapponicus* | *Calcarius lapponicus* | Lapland Longspur | Calcariidae | Passeriformes | |
| Bokje | *Lymnocryptes minimus* | *Lymnocryptes minimus* | Jack Snipe | Scolopacidae | Charadriiformes | |
| Grauwe kiekendief | *Circus pygargus* | *Circus pygargus* | Montagu's Harrier | Accipitridae | Accipitriformes | 14.17674152 |
| Valkparkiet | *Nymphicus hollandicus* | *Nymphicus hollandicus* | Cockatiel | Cacatuidae | Psittaciformes | |
| Amerikaanse smient | *Anas americana* | *Mareca americana* | American Wigeon | Anatidae | Anseriformes | |
| Witbuikrotgans | *Branta hrota* | *Branta bernicla* | Brant | Anatidae | Anseriformes | |
| Sneeuwgans | *Anser caerulescens* | *Anser caerulescens* | Snow Goose | Anatidae | Anseriformes | |
| Bijeneter | *Merops apiaster* | *Merops apiaster* | European Bee-eater | Meropidae | Coraciiformes | |
| Grasparkiet | *Melopsittacus undulatus* | *Melopsittacus undulatus* | Budgerigar | Psittaculidae | Psittaciformes | |
| Huiskraai | *Corvus splendens* | *Corvus splendens* | House Crow | Corvidae | Passeriformes | |

| Dutch Name | Scientific name (NDFF) | Scientific name (e_Bird) | English name | Family2 | Order2 | Home range radius (km) |
|---|---|---|---|---|---|---|
| Zwarte wouw | *Milvus migrans* | *Milvus migrans* | Black Kite | Accipitridae | Accipitriformes | |
| Kwak | *Nycticorax nycticorax* | *Nycticorax nycticorax* | Black-crowned Night-Heron | Ardeidae | Pelecaniformes | |
| Draaihals | *Jynx torquilla* | *Jynx torquilla* | Eurasian Wryneck | Picidae | Piciformes | 1.018871925 |
| Alk | *Alca torda* | *Alca torda* | Razorbill | Alcidae | Charadriiformes | |
| Siberische tjiftjaf | *Phylloscopus tristis* | *Phylloscopus collybita* | Common Chiffchaff | Phylloscopidae | Passeriformes | |
| Dwerggans | *Anser erythropus* | *Anser erythropus* | Lesser White-fronted Goose | Anatidae | Anseriformes | |
| Grauwe pijlstormvogel | *Puffinus griseus* | *Ardenna grisea* | Sooty Shearwater | Procellariidae | Procellariiformes | |
| Noordse stormvogel | *Fulmarus glacialis* | *Fulmarus glacialis* | Northern Fulmar | Procellariidae | Procellariiformes | |
| Middelste jager | *Stercorarius pomarinus* | *Stercorarius pomarinus* | Pomarine Jaeger | Stercorariidae | Charadriiformes | |
| Nachtzwaluw | *Caprimulgus europaeus* | *Caprimulgus europaeus* | European Nightjar | Caprimulgidae | Caprimulgiformes | 0.886002257 |
| Siberische boompieper | *Anthus hodgsoni* | *Anthus hodgsoni* | Olive-backed Pipit | Motacillidae | Passeriformes | |
| Bonte kraai | *Corvus cornix* | *Corvus cornix* | Hooded Crow | Corvidae | Passeriformes | |
| Kortsnavelboomkruiper | *Certhia familiaris macrodactyla* | *Certhia familiaris* | Eurasian Treecreeper | Certhiidae | Passeriformes | 0.216794834 |
| Grote burgemeester | *Larus hyperboreus* | *Larus hyperboreus* | Glaucous Gull | Laridae | Charadriiformes | |
| Grote karekiet | *Acrocephalus arundinaceus* | *Acrocephalus arundinaceus* | Great Reed Warbler | Acrocephalidae | Passeriformes | |
| Grote kruisbek | *Loxia pytyopsittacus* | *Loxia pytyopsittacus* | Parrot Crossbill | Fringillidae | Passeriformes | |
| Atlantische Canadese gans | *Branta canadensis canadensis* | *Branta canadensis* | Canada Goose | Anatidae | Anseriformes | |
| Pennantrosella | *Platycercus elegans* | *Platycercus elegans* | Crimson Rosella | Psittaculidae | Psittaciformes | |
| Kleine burgemeester | *Larus glaucoides* | *Larus glaucoides* | Iceland Gull | Laridae | Charadriiformes | |
| Bruine boszanger | *Phylloscopus fuscatus* | *Phylloscopus fuscatus* | Dusky Warbler | Phylloscopidae | Passeriformes | |
| Pallas' Boszanger | *Phylloscopus proregulus* | *Phylloscopus proregulus* | Pallas's Leaf Warbler | Phylloscopidae | Passeriformes | |
| Kleine vliegenvanger | *Ficedula parva* | *Ficedula parva* | Red-breasted Flycatcher | Muscicapidae | Passeriformes | |
| Dwerggors | *Emberiza pusilla* | *Emberiza pusilla* | Little Bunting | Emberizidae | Passeriformes | |
| Humes bladkoning | *Phylloscopus humei* | *Phylloscopus humei* | Hume's Leaf Warbler | Phylloscopidae | Passeriformes | |
| Hop | *Upupa epops* | *Upupa epops* | Eurasian Hoopoe | Upupidae | Bucerotiformes | 3.544009029 |
| Bergfluiter | *Phylloscopus bonelli* | *Phylloscopus bonelli* | Western Bonelli's Warbler | Phylloscopidae | Passeriformes | 0.187082869 |

| Dutch Name | Scientific name (NDFF) | Scientific name (e_Bird) | English name | Family2 | Order2 | Home range radius (km) |
|---|---|---|---|---|---|---|
| Pestvogel | *Bombycilla garrulus* | *Bombycilla garrulus* | Bohemian Waxwing | Bombycillidae | Passeriformes | |
| Witoogeend | *Aythya nyroca* | *Aythya nyroca* | Ferruginous Duck | Anatidae | Anseriformes | |
| Orpheusspotvogel | *Hippolais polyglotta* | *Hippolais polyglotta* | Melodious Warbler | Acrocephalidae | Passeriformes | 0.173205081 |
| Monniksparkiet | *Myiopsitta monachus* | *Myiopsitta monachus* | Monk Parakeet | Psittacidae | Psittaciformes | |
| Kleinste Canadese gans | *Branta hutchinsii minima* | *Branta hutchinsii* | Cackling Goose | Anatidae | Anseriformes | |
| Fluiter | *Phylloscopus sibilatrix* | *Phylloscopus sibilatrix* | Wood Warbler | Phylloscopidae | Passeriformes | |
| Grauwe gors | *Emberiza calandra* | *Emberiza calandra* | Corn Bunting | Emberizidae | Passeriformes | |
| Kuifleeuwerik | *Galerida cristata* | *Galerida cristata* | Crested Lark | Alaudidae | Passeriformes | |
| Kanarie | *Serinus canaria* | *Serinus canaria* | Atlantic Canary | Fringillidae | Passeriformes | |
| Noordse kauw | *Corvus monedula monedula* | *Corvus monedula* | Eurasian Jackdaw | Corvidae | Passeriformes | |
| Kaapse casarca | *Tadorna cana* | *Tadorna cana* | South African Shelduck | Anatidae | Anseriformes | |
| Woudaap | *Ixobrychus minutus* | *Ixobrychus minutus* | Little Bittern | Ardeidae | Pelecaniformes | |
| Grote grijze snip | *Limnodromus scolopaceus* | *Limnodromus scolopaceus* | Long-billed Dowitcher | Scolopacidae | Charadriiformes | |
| Taigaboomkruiper | *Certhia familiaris* | *Certhia familiaris* | Eurasian Treecreeper | Certhiidae | Passeriformes | 0.216794834 |
| Arendbuizerd | *Buteo rufinus* | *Buteo rufinus* | Long-legged Buzzard | Accipitridae | Accipitriformes | |
| Sneeuwuil | *Bubo scandiacus* | *Bubo scandiacus* | Snowy Owl | Strigidae | Strigiformes | |
| Slangenarend | *Circaetus gallicus* | *Circaetus gallicus* | Short-toed Snake-Eagle | Accipitridae | Accipitriformes | 8.860022573 |
| Sperweruil | *Surnia ulula* | *Surnia ulula* | Northern Hawk Owl | Strigidae | Strigiformes | |
| Witwangstern | *Chlidonias hybrida* | *Chlidonias hybrida* | Whiskered Tern | Laridae | Charadriiformes | |
| Zebravink | *Taeniopygia guttata* | *Taeniopygia guttata* | Zebra Finch | Estrildidae | Passeriformes | |
| Dwerguil | *Glaucidium passerinum* | *Glaucidium passerinum* | Eurasian Pygmy-Owl | Strigidae | Strigiformes | 1.118033989 |
| Brilzee-eend | *Melanitta perspicillata* | *Melanitta perspicillata* | Surf Scoter | Anatidae | Anseriformes | |
| Witbandkruisbek | *Loxia leucoptera* | *Loxia leucoptera* | Two-barred Crossbill | Fringillidae | Passeriformes | |
| Siberische taling | *Sibirionetta formosa* | *Sibirionetta formosa* | Baikal Teal | Anatidae | Anseriformes | |
| Rood kamhoen | *Gallus gallus* | *Gallus gallus* | Red Junglefowl | Phasianidae | Galliformes | |
| Korhoen | *Tetrao tetrix* | *Lyrurus tetrix* | Black Grouse | Phasianidae | Galliformes | |
| Swinhoes boszanger | *Swinhoes boszanger* | *Phylloscopus plumbeitarsus* | Two-barred Warbler | Phylloscopidae | Passeriformes | |
| Kleine Canadese gans | *Branta hutchinsii* | *Branta hutchinsii* | Cackling Goose | Anatidae | Anseriformes | |
| Kokardezaagbek | *Lophodytes* | *Lophodytes* | Hooded | Anatidae | Anseriformes | |

| Dutch Name | Scientific name (NDFF) | Scientific name (e_Bird) | English name | Family2 | Order2 | Home range radius (km) |
|---|---|---|---|---|---|---|
| | *cucullatus* | *cucullatus* | Merganser | | | |
| Manengans | *Chenonetta jubata* | *Chenonetta jubata* | Maned Duck | Anatidae | Anseriformes | |
| Caribische flamingo | *Phoenicopterus ruber* | *Phoenicopterus ruber* | American Flamingo | Phoenicopteridae | Phoenicopteriformes | |
| Bruine klauwier | *Lanius cristatus* | *Lanius cristatus* | Brown Shrike | Laniidae | Passeriformes | |
| Bahamapijlstaart | *Anas bahamensis* | *Anas bahamensis* | White-cheeked Pintail | Anatidae | Anseriformes | |
| Notenkraker | *Nucifraga caryocatactes* | *Nucifraga caryocatactes* | Spotted Nutcracker | Corvidae | Passeriformes | 0.363774656 |
| Kroonkraanvogel | *Balearica pavonina* | *Balearica pavonina* | Black Crowned-Crane | Gruidae | Gruiformes | |
| Geelsnavelduiker | *Gavia adamsii* | *Gavia adamsii* | Yellow-billed Loon | Gaviidae | Gaviiformes | |
| Ringtaling | *Callonetta leucophrys* | *Callonetta leucophrys* | Ringed Teal | Anatidae | Anseriformes | |
| Ringsnaveleend | *Aythya collaris* | *Aythya collaris* | Ring-necked Duck | Anatidae | Anseriformes | |
| Geronticus eremita | *Geronticus eremita* | *Geronticus eremita* | Northern Bald Ibis | Threskiornithidae | Pelecaniformes | |
| Noordse goudvink | *Pyrrhula pyrrhula pyrrhula* | *Pyrrhula pyrrhula* | Eurasian Bullfinch | Fringillidae | Passeriformes | |
| Grote aalscholver | *Phalacrocorax carbo carbo* | *Phalacrocorax carbo* | Great Cormorant | Phalacrocoracidae | Suliformes | |
| Kleine trap | *Tetrax tetrax* | *Tetrax tetrax* | Little Bustard | Otididae | Otidiformes | |
| Heilige ibis | *Threskiornis aethiopicus* | *Threskiornis aethiopicus* | African Sacred Ibis | Threskiornithidae | Pelecaniformes | |
| Blauwe pauw | *Pavo cristatus* | *Pavo cristatus* | Indian Peafowl | Phasianidae | Galliformes | |
| Kleine alk | *Alle alle* | *Alle alle* | Little Auk | Alcidae | Charadriiformes | |
| Helmparelhoen | *Numida meleagris* | *Numida meleagris* | Helmeted Guineafowl | Numididae | Galliformes | |
| Japanse nachtegaal | *Leiothrix lutea* | *Leiothrix lutea* | Red-billed Leiothrix | Leiothrichidae | Passeriformes | |
| Bruinkopdiksnavelmees | *Sinosuthora webbiana* | *Sinosuthora webbiana* | Vinous-throated Parrotbill | Sylviidae | Passeriformes | |
| Geelvleugelara | *Ara macao* | *Ara macao* | Scarlet Macaw | Psittacidae | Psittaciformes | |
| Goudfazant | *Chrysolophus pictus* | *Chrysolophus pictus* | Golden Pheasant | Phasianidae | Galliformes | |
| Ross' Gans | *Anser rossii* | *Anser rossii* | Ross’s Goose | Anatidae | Anseriformes | |
| Ortolaan | *Emberiza hortulana* | *Emberiza hortulana* | Ortolan Bunting | Emberizidae | Passeriformes | |
| Magelhaengans | *Chloephaga picta* | *Chloephaga picta* | Upland Goose | Anatidae | Anseriformes | |
| Kleine flamingo | *Phoeniconaias minor* | *Phoeniconaias minor* | Lesser Flamingo | Phoenicopteridae | Phoenicopteriformes | |
| Iberische tjiftjaf | *Phylloscopus ibericus* | *Phylloscopus ibericus* | Iberian Chiffchaff | Phylloscopidae | Passeriformes | |
| Kaneeltaling | *Anas cyanoptera* | *Spatula cyanoptera* | Cinnamon Teal | Anatidae | Anseriformes | |
| Duinpieper | *Anthus campestris* | *Anthus campestris* | Tawny Pipit | Motacillidae | Passeriformes | |

| Dutch Name | Scientific name (NDFF) | Scientific name (e_Bird) | English name | Family2 | Order2 | Home range radius (km) |
|---|---|---|---|---|---|---|
| Kwartelkoning | *Crex crex* | *Crex crex* | Corn Crake | Rallidae | Gruiformes | 0.207364414 |
| Noordse nachtegaal | *Luscinia luscinia* | *Luscinia luscinia* | Thrush Nightingale | Muscicapidae | Passeriformes | |
| Vlekbekeend | *Anas poecilorhyncha* | *Anas poecilorhyncha* | Indian Spot-billed Duck | Anatidae | Anseriformes | |
| Krekelzanger | *Locustella fluviatilis* | *Locustella fluviatilis* | River Warbler | Locustellidae | Passeriformes | |
| Lady-amherstfazant | *Chrysolophus amherstiae* | *Chrysolophus amherstiae* | Lady Amherst's Pheasant | Phasianidae | Galliformes | |
| Paradijscasarca | *Tadorna variegata* | *Tadorna variegata* | Paradise Shelduck | Anatidae | Anseriformes | |
| Grauwe fitis | *Phylloscopus trochiloides* | *Phylloscopus trochiloides* | Greenish Warbler | Phylloscopidae | Passeriformes | |
| Bronskopeend | *Anas falcata* | *Mareca falcata* | Falcated Duck | Anatidae | Anseriformes | |
| Oevermaina | *Acridotheres ginginianus* | *Acridotheres ginginianus* | Bank Myna | Sturnidae | Passeriformes | |
| Kalkoen | *Meleagris gallopavo* | *Meleagris gallopavo* | Wild Turkey | Phasianidae | Galliformes | |
| Marmereend | *Marmaronetta angustirostris* | *Marmaronetta angustirostris* | Marbled Duck | Anatidae | Anseriformes | |
| Toendrarietgans | *Anser serrirostris* | *Anser serrirostris* | Tundra Bean-Goose | Anatidae | Anseriformes | |
| Jufferkraanvogel | *Grus virgo* | *Anthropoides virgo* | Demoiselle Crane | Gruidae | Gruiformes | |
| Papegaaiduiker | *Fratercula arctica* | *Fratercula arctica* | Atlantic Puffin | Alcidae | Charadriiformes | |
| Trompetzwaan | *Cygnus buccinator* | *Cygnus buccinator* | Trumpeter Swan | Anatidae | Anseriformes | |
| Witkruintapuit | *Oenanthe leucopyga* | *Oenanthe leucopyga* | White-crowned Wheatear | Muscicapidae | Passeriformes | |
| Hawaigans | *Branta sandvicensis* | *Branta sandvicensis* | Nene | Anatidae | Anseriformes | |
| Kleine topper | *Aythya affinis* | *Aythya affinis* | Lesser Scaup | Anatidae | Anseriformes | |
| Groenlandse kolgans | *Anser albifrons flavirostris* | *Anser albifrons* | Greater White-fronted Goose | Anatidae | Anseriformes | |
| Oosterse tortel | *Streptopelia orientalis* | *Streptopelia orientalis* | Oriental Turtle-Dove | Columbidae | Columbiformes | |
| Bont boertje | *Poicephalus senegalus* | *Poicephalus senegalus* | Senegal Parrot | Psittacidae | Psittaciformes | |
| Taigarietgans | *Anser fabalis* | *Anser fabalis* | Taiga Bean-Goose | Anatidae | Anseriformes | |
| Koningsfazant | *Syrmaticus reevesii* | *Syrmaticus reevesii* | Reeves's Pheasant | Phasianidae | Galliformes | |
| Graszanger | *Cisticola juncidis* | *Cisticola juncidis* | Zitting Cisticola | Cisticolidae | Passeriformes | 0.12 |
| Zwaangans | *Anser cygnoides* | *Anser cygnoides* | Swan Goose | Anatidae | Anseriformes | |
| Mexicaanse roodmus | *Haemorhous mexicanus* | *Haemorhous mexicanus* | House Finch | Fringillidae | Passeriformes | |
| Woestijntapuit | *Oenanthe deserti* | *Oenanthe deserti* | Desert Wheatear | Muscicapidae | Passeriformes | |
| Grijze junco | *Junco hyemalis* | *Junco* | Dark-eyed | Passerellidae | Passeriformes | |

| Dutch Name | Scientific name (NDFF) | Scientific name (e_Bird) | English name | Family2 | Order2 | Home range radius (km) |
|---|---|---|---|---|---|---|
| | | *hyemalis* | Junco | | | |
| Amerikaanse oeverloper | *Actitis macularius* | *Actitis macularius* | Spotted Sandpiper | Scolopacidae | Charadriiformes | |
| Roodsterblauwborst | *Roodsterblauwborst* | *Luscinia svecica* | Bluethroat | Muscicapidae | Passeriformes | |
| Lachduif | *Streptopelia risoria* | *Streptopelia roseogrisea* | African Collared-Dove | Columbidae | Columbiformes | |
| Blauwvleugelgans | *Cyanochen cyanoptera* | *Cyanochen cyanoptera* | Blue-winged Goose | Anatidae | Anseriformes | |
| Ruigpootuil | *Aegolius funereus* | *Aegolius funereus* | Boreal Owl | Strigidae | Strigiformes | 1.772004515 |
| Australische bergeend | *Tadorna tadornoides* | *Tadorna tadornoides* | Australian Shelduck | Anatidae | Anseriformes | |
| Rode kardinaal | *Cardinalis cardinalis* | *Cardinalis cardinalis* | Northern Cardinal | Cardinalidae | Passeriformes | |
| Stormvogeltje | *Hydrobates pelagicus* | *Hydrobates pelagicus* | European Storm-Petrel | Hydrobatidae | Procellariiformes | |
| Roodkeelnachtegaal | *Calliope calliope* | *Calliope calliope* | Siberian Rubythroat | Muscicapidae | Passeriformes | |
| Peposacaeend | *Netta peposaca* | *Netta peposaca* | Rosy-billed Pochard | Anatidae | Anseriformes | |
| Zwartbuikwaterspreeuw | *Cinclus cinclus cinclus* | *Cinclus cinclus* | White-throated Dipper | Cinclidae | Passeriformes | |
| Kumliens meeuw | *Larus glaucoides kumlieni* | *Larus glaucoides* | Iceland Gull | Laridae | Charadriiformes | |

*Note*: Columns include Dutch common name, scientific name according to the NDFF dataset NDFFdataset, https://ndff.nl/ ; "Scientific name (NDFF)") and the eBird dataset https://ebird.org/home ; "Scientific name (e_Bird)"), English common name, taxonomic family (Family2), taxonomic order (Order2), and home-range radius (km) used in the study analyses (Tamburello et al., 2015). Home-range radius values represent species' estimated home-range size. Of the 374 bird species considered in the Netherlands, 50 species could be matched to the home-range dataset; these 50 species were used as a reference to guide the selection of analysis spatial scale (grid size). Because 49% of the matched species had home-range radii < 200 m, we set the primary analysis grid size to 200 m and used 100 m and 300 m as sensitivity analyses.

**Table S3**: Pairwise Pearson correlation coefficients (r)

| | **Habitat heterogeneity** | **Vegetation heterogeneity** | **Foliage layer richness** |
|---|---|---|---|
| **Habitat heterogeneity** | 1.0000000 | 0.1630553 | 0.3926048 |
| **Vegetation heterogeneity** | 0.1630553 | 1.0000000 | 0.1945106 |
| **Foliage layer richness** | 0.3926048 | 0.1945106 | 1.0000000 |

*Note:* Pairwise pearson correlation coefficients among environmental heterogeneity predictors: plant species richness, habitat heterogeneity and foliage-layer richness. Values in the cells indicate correlation strength; colours show direction and magnitude (red = positive, blue = negative). All correlations are weak to moderate (|r| ≤ 0.4), indicating limited collinearity among predictors.

**Table S4**: Variance Inflation Factors (VIF)

| **Predictor** | **VIF** |
|---|---|
| Habitat heterogeneity | 1.19 |
| Vegetation heterogeneity | 1.05 |
| Foliage layer richness | 1.21 |

*Note*: All VIF values were below 2, indicating no evidence of problematic multicollinearity.

**Table S5**: Basis dimension (k) checking results for GAM (200 m scale; REML)

| Smooth term | *k'* | *edf* | *k*-index | *p*-value |
|---|---|---|---|---|
| *s*(HH) | 9.00 | 3.48 | 0.99 | 0.17 |
| *s*(PlantSR) | 9.00 | 5.73 | 1.02 | 0.94 |
| *s*(FR) | 9.00 | 7.85 | 1.00 | 0.46 |
| *ti*(VH):Built | 4.00 | 3.24 | 1.02 | 0.95 |
| *ti*(HH_200m):Built | 4.00 | 3.22 | 0.99 | 0.14 |
| *ti*(foliage_richness):Built | 4.00 | 3.91 | 1.00 | 0.45 |

**Table S6**: Concurvity diagnostics for the GAM (200 m scale)

| Model term | Worst | Observed | Estimate |
|---|---|---|---|
| Parametric component (“para”) | 0.362 | 0.362 | 0.362 |
| *s*(HH_200m) | 0.720 | 0.619 | 0.555 |
| *s*(PlantSR_200m) | 0.635 | 0.573 | 0.563 |
| *s*(FR) | 0.716 | 0.640 | 0.580 |

| Model term | Worst | Observed | Estimate |
|---|---|---|---|
| *ti*(VH_200m, built) | 0.635 | 0.568 | 0.551 |
| *ti*(HH_200m, built) | 0.726 | 0.700 | 0.560 |
| *ti*(FR, built) | 0.729 | 0.611 | 0.622 |

Note: Concurvity was generally moderate (worst-case 0.63–0.73 for smooth and interaction terms), indicating no severe redundancy among smooth terms.

**Table S7**: Comparison of GAM performance metrics across spatial scales (100 m, 200 m, 300 m)

| **scale_m** | **n** | ***adjR2*** | **dev_exp (%)** |
|---|---|---|---|
| 100 | 237889 | 0.132 | 13.2 |
| 200 | 66337 | 0.211 | 21.1 |
| 300 | 30666 | 0.259 | 26 |

**Table S8**: Generalized Additive Model (GAM) statistics for the effects of environmental heterogeneity on bird richness across urbanization intensity (100m)

| *Predictors* | *edf* | *Ref.df* | *F* | *p-value* |
|---|---|---|---|---|
| Habitat heterogeneity | 7.30 | 9 | 39.21 | <0.001 |
| Habitat heterogeneity × Built-up area | 3.56 | 4 | 51.98 | <0.001 |
| Vegetation heterogeneity | 6.25 | 9 | 2224.98 | <0.001 |
| Vegetation heterogeneity × Built-up area | 3.68 | 4 | 295.31 | <0.001 |
| Foliage richness | 7.29 | 9 | 17.35 | <0.001 |
| Foliage richness × Built-up area | 3.95 | 4 | 130.97 | <0.001 |

**Table S9**: Generalized Additive Model (GAM) statistics for the effects of environmental heterogeneity on bird richness across urbanization intensity (300m)

| Predictors | *edf* | *Ref.df* | *F* | *p-value* |
|---|---|---|---|---|
| Plant richness | 5.61 | 9 | 546.39 | <0.001 |
| Plant richness × Built-up area | 3.25 | 4 | 40.25 | <0.001 |
| Foliage richness | 7.81 | 9 | 12.98 | <0.001 |
| Foliage richness × Built-up area | 3.81 | 4 | 19.70 | <0.001 |
| Habitat heterogeneity | 1.39 | 9 | 12.14 | <0.001 |
| Habitat heterogeneity × Built-up area | 3.21 | 4 | 14.30 | <0.001 |

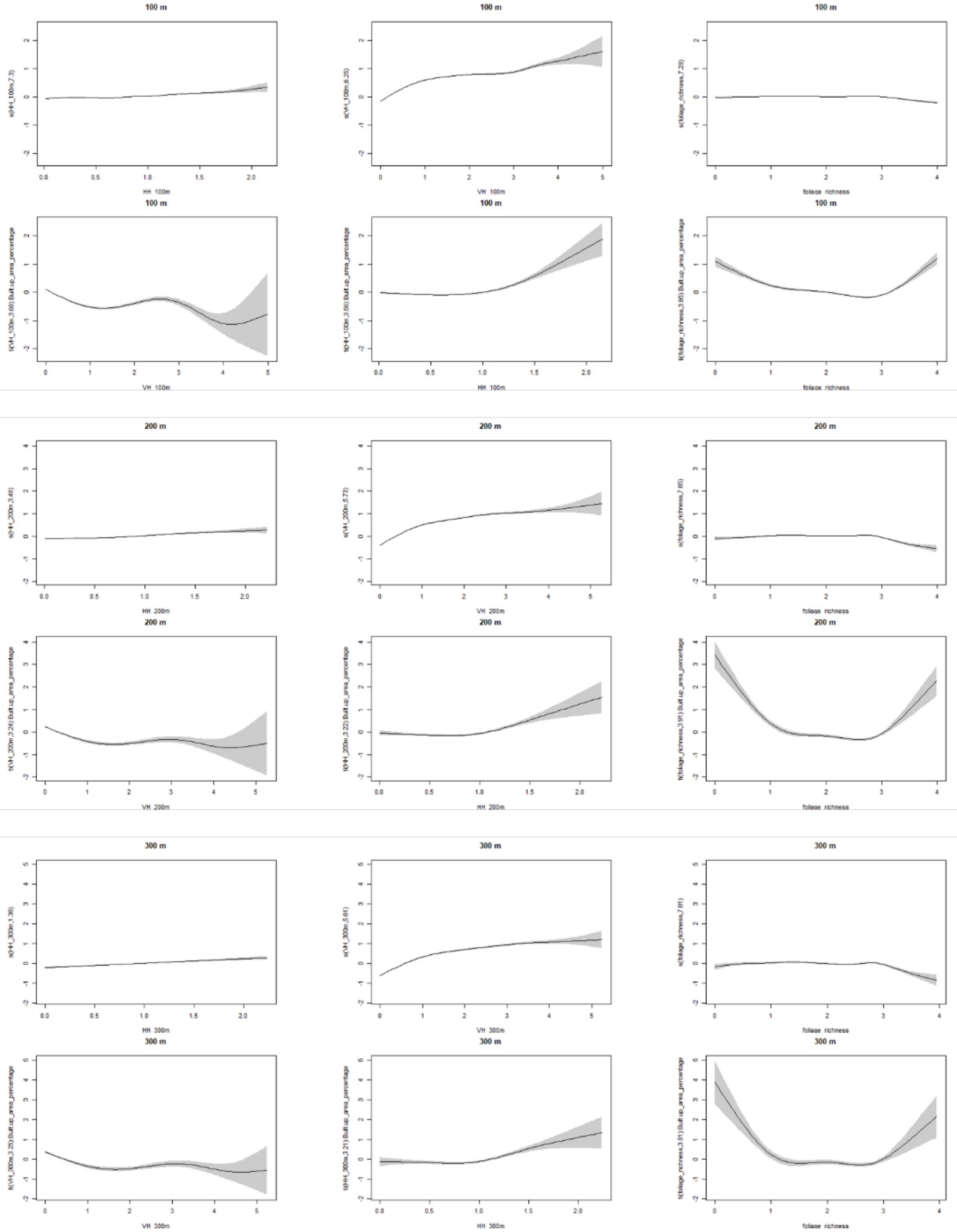


**Figure S2**: Scale sensitivity of GAM-estimated effects for habitat heterogeneity, vegetation height heterogeneity, plant richness, and their interactions with built-up area.

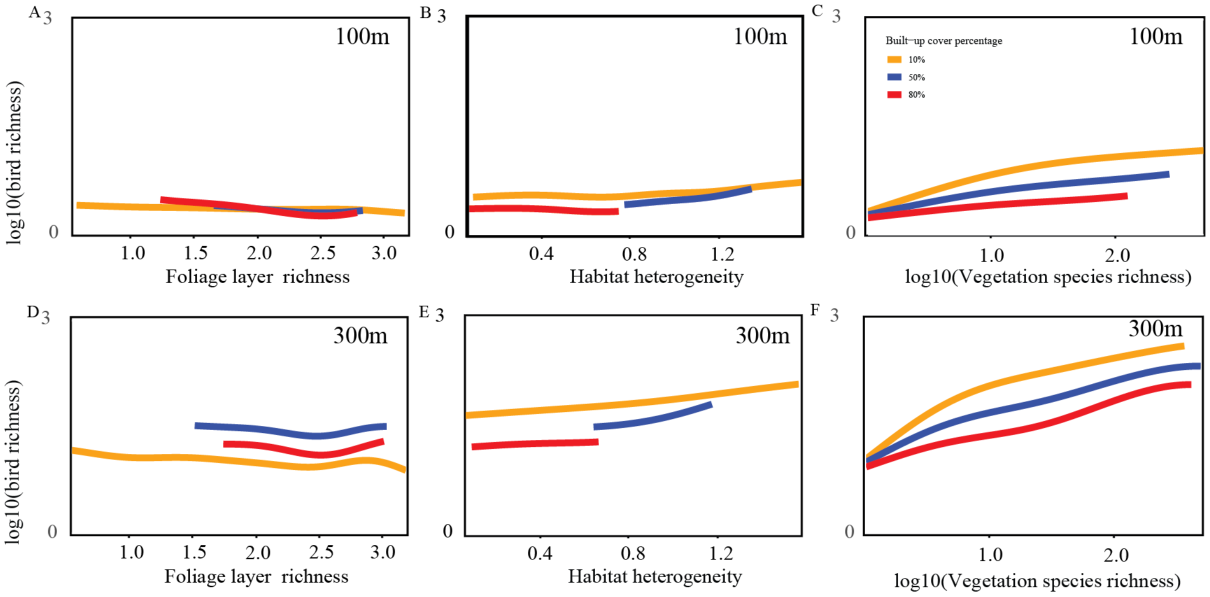


**Figure S3**: Environmental heterogeneity effects on urban bird richness across urbanisation intensity and scale. Conditional partial response curves from GAMs showing relationships between bird richness and (A, D) foliage layer richness, (B, E) habitat heterogeneity, and (C, F) plant species richness at two spatial scales: 100 m (A–C) and 300 m (D–F). Coloured lines show model predictions at three representative built-up cover levels (10%, 50%, 80. Bird richness was modelled as log(1 + richness).

Tamburello, N., Côté, I. M., & Dulvy, N. K. (2015). Energy and the scaling of animal space use. *The American Naturalist*, *186*(2), 196-211.
